# Determining gene specificity from multivariate single-cell RNA sequencing data

**DOI:** 10.1101/2025.11.21.689845

**Authors:** Nikhila P. Swarna, A. Sina Booeshaghi, Elisabeth Rebboah, M. Grace Gordon, Pooja Kathail, Taibo Li, Marcus Alvarez, Chun Jimmie Ye, Barbara Wold, Ali Mortazavi, Lior Pachter

## Abstract

**Background:** An important application of single-cell genomics experiments is to identify genes specific to biological categories or experimental conditions. We develop ember (Entropy Metrics for Biological ExploRation), an entropy-based framework that uses Shannon’s chain rule across nested biological partitions. Four axioms characterize its Ψ score within a class of normalized entropy-decomposition scores.

**Results:** Applying ember to eight tissues from eight founder mouse strains, we find that gene specificity is often unintuitive: canonical markers can be supplanted, housekeeping genes are context-dependent, and mouse strain can drive unexpected cell type switching. Unsupervised learning on entropy metrics uncovers shared genes specialized to male gonads and kidney, as well as genes specific to non-consecutive developmental stages in the kidney. To facilitate further exploration of gene specificity in mice, we have also developed a comprehensive specificity database, along with a web interface, API and MCP server. Extending ember to a human PBMC dataset collected from 255 diverse individuals, we find that variation in PBMCs is largely localized to classical monocytes. We also find genes with unique specificity by sex, age and ancestral background.

**Conclusions:** Together, these applications establish ember as a powerful tool and provide a roadmap for elucidating the impact of human genetic variation using the murine model.

## Background

With the advent of combinatorial barcoding and split-pool technologies in single-cell RNA sequencing (scRNA-seq), experiments have become increasingly high-dimensional (1, 2). Modern experiments not only assay tens of thousands of genes, but can also accommodate replicates across multiple biological variables: sex, tissue, age, genotype, and cell states (3–8). Although this expansion in experimental size paired with an increased depth of sequencing allows unprecedented resolution of cellular and molecular biology, there is a lack of tools to appropriately dissect datasets and extract meaningful insights (9–11).

A fundamental challenge in analyzing such multivariate data is identifying genes that are highly specific to a given biological variable. Gene specificity underlies marker discovery, cell-type annotation, interpretation of developmental programs, and identification of condition-dependent molecular phenotypes, making it a central concept in single-cell analysis (12, 13). Several measures of specificity have been proposed to address this problem. For example, Tau, or the tissue specificity index (TSI), summarizes gene specificity as a single number bounded between 0 and 1, with 0 indicating broad expression and 1 indicating perfect specificity (14, 15). The preferential expression measure (PEM) assigns positive or negative values indicating over- or under-expression in a given tissue (16, 17), producing tissue-level scores that can subsequently be aggregated. The Gini index, originally developed in economics to quantify inequality, has also been adapted to describe the unevenness of gene expression across tissues (18–20). Although these specificity indices have been compared empirically (21), such studies typically evaluate performance by the number of genes identified as specific rather than by examining the mathematical properties that define specificity itself.

Measures such as Tau, PEM and Gini summarize different aspects of expression imbalance. Our aim is to track the single-cell expression distribution through predefined, nested biological partitions using an entropy decomposition. Generalized linear model frameworks such as DESeq2 (22) and mixed-model approaches such as variancePartition (23) provide decomposable models but rely on parametric assumptions and can scale poorly to increasingly large datasets. Moreover, these methods were originally developed for bulk expression data and therefore typically require pseudobulk-level summaries when applied to single-cell experiments, discarding valuable information about how expression is distributed across individual cells.

Our normalized within-block entropy score is bounded between 0 and 1 when total entropy is positive, is insensitive to category labels, and obeys a precise refinement identity inherited from Shannon’s chain rule (*Supplementary Note*). Four axioms specify regularity, symmetry, hierarchical grouping of the underlying information functional, and normalization by its total value. Faddeev’s theorem characterizes that functional up to a multiplicative constant, which cancels in this score, the uniqueness result is within this class of entropy-decomposition scores (24, 25). Casey *et al*. also give an additive decomposition of single-cell expression heterogeneity into between- and within-cluster components, their construction can be iterated over nested partitions (26). Their quantity is Kullback–Leibler divergence from uniform expression per cell, so its between-cluster term compares a cluster’s expression share with its share of cells. Here we normalize the within-block contribution of Shannon entropy and use it, together with block scores, to analyze predefined multivariate biological partitions.

We introduce three complementary entropy metrics: Ψ (Psi): fraction of gene-expression entropy retained within partition blocks, *ψ*_*block*_ (psi_block): specificity to a block, and *ζ* (Zeta): specificity to a partition. Detailed derivations and interpretations are provided in the *Methods* section. Under these four axioms, Ψ is uniquely determined within the normalized entropy-decomposition construction and is independent of the multiplicative normalization of Shannon entropy (*Supplementary Note*). Our framework is implemented in the open-source Python package ember (Entropy Metrics for Biological ExploRation).

To showcase the novelty and utility of our framework, we benchmark ember against existing specificity measures (Tau, PEM, Gini, Deseq2, and variancePartition) across simulated expression patterns (*Supplementary Note*). While these methods recover similar results for simple single-block specificity, they differ for more complex patterns: Tau and Gini struggle with multiblock and multivariate specificity, DESeq2 and variancePartition do not consistently distinguish differential or variable expression from specific expression, and PEM performs similarly to ember after aggregation but cannot distinguish expression patterns with identical pseudobulk profiles that differ at single-cell resolution. These comparisons define the settings in which ember is useful and the behavior that follows from the assumptions of our framework.

Together with novel visualization strategies, ember provides a principled and interpretable toolkit for studying biological specificity in high-dimensional single-cell data. Rather than centering the main text on methodological comparisons, we focus on the biological questions that this framework enables. We use three large multivariate single-cell datasets: (1) snRNA-seq of eight tissues from eight individuals across the eight founder mouse strains (8cube founder data (6)), (2) scRNA-seq of murine kidney across six developmental time points (developmental kidney data (7)), and (3) snRNA-seq of human PBMCs from 255 diverse individuals (human PBMC data). Through a collection of case studies from these data, we demonstrate how information-theoretic measures of specificity can both zoom-in to identify genes exhibiting complex multivariate patterns of condition specificity and zoom-out to reveal higher-order principles of biological organization across cell types, developmental stages and genetic backgrounds.

## Results

### Gene specificity is unintuitive and hierarchical decomposition allows for novel gene discovery

#### Strain-specific genes

Applying ember to the 8cube founder data set (6), we identified genes that are strain-specific across all eight tissues (Fig. 1a). We identified *Cwc22*, which encodes spliceosome-associated protein 58, as a WSBEiJ (WSBJ) specific gene. *Cwc22* has been documented to have heightened expression in WSBJ mice due to a copy number expansion at the *R2d2* locus, located approximately 6 Mb downstream from its paralog *R2d1* (30) (Supp. Fig 1c). We also identified the gap junction protein encoded by *Gjb4*, as specific to NZO/HlLtJ (NZOJ) (Supp Fig 1d). It has been documented that *Gjb4* expression in diabetic NZOJ mice is enhanced by a reduced *miR-341-3p* expression (31).

We identified *Tas1r1* as a CASTEiJ (CASTJ) specific gene, with a slight bias towards females (Supp Fig 1a). *Tas1r1* encodes a member of the taste receptor family of genes, and functions as a receptor of umami taste along with another member of the same family, *Tas1r3* (32). Interestingly, it has also been observed that this wild-derived strain displays a unique oral preference to sugars, distinguishing CASTJ from other strains of mice and rats (33). When comparing transcriptomic differences between mouse strains mapped to the standard GRCm39 reference genome, we run the risk of falsely quantifying mapping artifacts as strain specificity becauase the GRCm39 assembly is derived from a C57BL/6J (B6J) background (6). To exclude the possibility that *Tas1r1*’s specificity to CASTJ mice could be the result of such a mapping artifact, we examined its coding sequence from the recent first release of the telomere-to-telomere (T2T) genomes for B6J and CASTJ (34, 35) (Supp. table 1). There is no evidence of frameshift or non-sense mutations and no alternative splicing, as the total CDS and protein length are conserved. This provides no evidence that *Tas1r1* is a pseudogene in B6J relative to CASTJ. Between the strains, 19 SNPs occur in exonic regions, of which three cause amino acid substitutions that may alter protein structure. We observed a large 4396 base pair deletion in B6J compared to CASTJ in the fourth intron at the CASTJ T2T coordinates 4:153181901-153186296 (reverse strand). Large intronic deletions have been associated with over or under expression of genes (36–38), thus cluing why we may be observing this specificity of *Tas1r1* to CASTJ. The gene just upstream of *Tas1r1* on the reverse strand chromosome four, *Zbtb48*, also exhibits biased expression towards CASTJ, though not as striking as that of *Tas1r1* (Supp Fig 1b), further validating our finding.

**Fig. 1.**
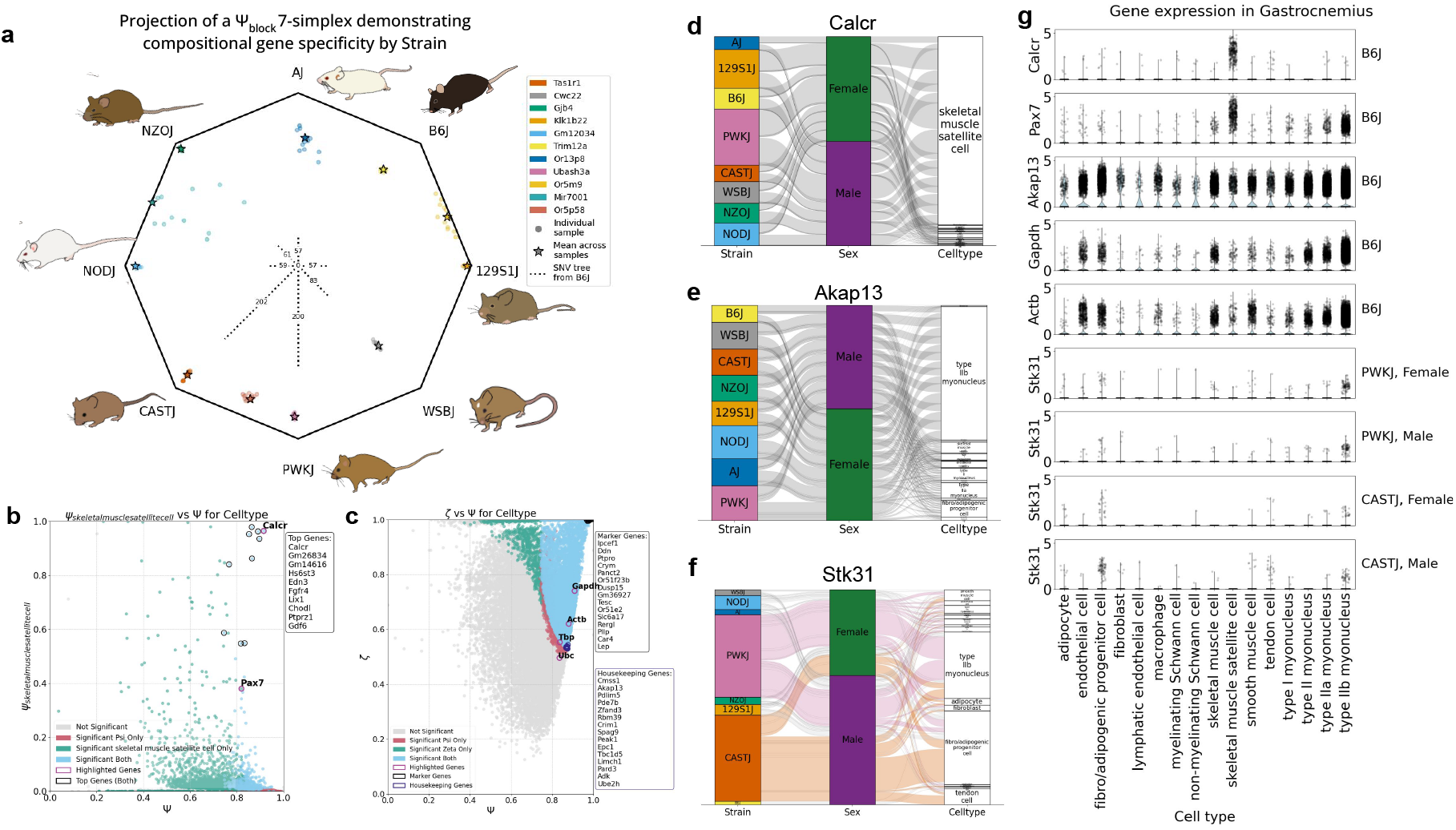
Visualizing gene specificity and finding highly specific and non-specific genes a. Octagonal plot showing gene specificity in compositional space. The 8cube founder dataset (6) was partitioned by strain to generate *ψ*_*block*_ scores for each of the eight founder strains. Each gene is represented as a coordinate on a 7-simplex, where each vertex indicates perfect specificity to one strain. We project genes according to single nucleotide variant distance to B6J in millions taken from Ferraj *et al. 2023* (27). Selected genes show specificity to one or two strains. Fifty balanced samples across biological replicates were used to generate *ψ*_*block*_ scores. Each sample is shown as a grey circle, and the Aitchison mean of these 50 samples is shown as a star. Mouse images from Saul *et al*. 2019 (28) **b**. Scatter plot showing gene specificity to skeletal muscle satellite cells in 8cube gastrocnemius (6), partitioned by cell type. Genes with high Ψ and high *ψ*_*skeletalmusclesatellitecells*_ are identified as marker genes (highlighted in black). Significance testing for both metrics is shown by color: genes significant for both are light blue and exhibit reliable specificity. Top marker genes are listed in the top-right text box. **c**. Scatter plot showing gene specificity in 8cube gastrocnemius (6), partitioned by cell type. Genes with high Ψ and high *ζ* are highly specific (highlighted in black), whereas those with high Ψ and low *ζ* are non-specific (highlighted in violet). Canonical housekeeping genes are highlighted in purple. Top cell type–specific genes are listed in the top-right text box, and top housekeeping genes are listed in the bottom-right text box. **d, e, f**. Alluvial plots depicting pseudo-bulked gene expression of *Calcr, Akap13* and *Stk31* across strains, sexes and cell types in 8cube gastrocnemius (6), generated using wompwomp (29). The gene *Stk31* was identified from gastrocnemius data by partitioning by strain and cell type and using resultant *ψ*_*block*_ scores to filter for the desired specificty pattern. **g**.Violin plots of gene expression in 8cube gastrocnemius (6), with gene names on the left and strain/sex annotations on the right.

In addition to strain-specific genes, we also observed genes that are specific to pairs of strains. *Or5m9* encodes an olfactory receptor specific to B6J and 129S1/SvImJ (129S1J). Another olfactory receptor *Or5p58* is specific to the wild-derived strains PWK/PhJ (PWKJ) and CASTJ, the most genetically distant from B6J of the eight founder strains. *Mir7001*, encodes a microRNA specific to the two diabetic mouse strains NZOJ and NOD/ShiLtJ (NODJ). Additionally, we found that gene specificity to mouse strain is variable across biological replicates. We applied a novel visualization strategy to depict *ψ*_*block*_ scores for each founder mouse strain and their mean and standard deviation values in compositional space as an octagon, where proximity to a vertex represents greater specificity (Fig. 1a, detailed explanation in *Methods*).

It should be noted that a gene that is biologically relevant need not be the most specific gene to a partition. For instance, *Cwc22* is well studied example of a strain-specific gene (30), however its specificity scores are *ζ* = 0.46 and Ψ = 0.90. By the definition of *ζ* (see *Methods*), a score of 0.46 which is closer to 0 than it is to 1 would suggest *Cwc22* is not a “highly-specific” gene to WSBJ mice. Examining its expression closely, we see reasonable expression in every mouse strain, which is why the *ζ* score is not closer to 1 (Supp Fig 1c). This does not however undercut the biological significance of *Cwc22*’s specific biology in WSBJ mice.

Specificity metrics are, by formulation, subjective and context dependent. A series of simulation benchmarking experiments showcase how the exact numerical values of a gene’s specificity can change with normalization strategy, cell depth and partition annotation (*Supplementary Note*). This makes biologically relevant thresholding of entropy metrics a meaningful strategy to ensure reproducible lists of context-specific genes across technical modalities. And using genes with biological validation to inform specificity thresholds within the context being studied (like *Cwc22* for strain specificity in founder mice) can improve interpretability. These results show that ember can identify known and novel genes that are specific to one or more mouse strains while accounting for biological context.

#### Cell type markers and housekeeping genes

Our method ember can be used to find cell type markers and housekeeping genes, while taking into account all possible partitions in a multivariate dataset. We found markers for skeletal muscle satellite cells, a rare but essential stem cell population required for adult muscle repair, making up only 1% of all nuclei in the 8cube gastrocnemius dataset (6) (Fig. 1d). The top satellite stem cell marker to emerge from our analysis was *Calcr*, calcitonin receptor (Fig. 1b, 1d). *Calcr* maintains the muscle stem cell pool by keeping them in a dormant state and in their location via the *cAMP-PKA* pathway (39). We were surprised that *Pax7*, the canonical myogenic marker, did not appear in our list of top satellite cell markers. In our analysis, *Pax7* was significant after permutation testing by *ψ*_*satellitecells*_ but not by Ψ (Fig 1b), which means that *Pax7* has some biased expression in satellite cells, but partitioning by cell type does not explain its expression pattern. Examining total expression across all gastrocnemius cell types, we found that *Pax7* also has significant expression in type IIB myoneuceli, the most populous of the gastrocnemius cell types (Fig. 1g). We validated these findings by comparing to a gastrocnemius single-nucleus dataset generated by Alexander S. Ham *et al*. 2025 (40), where we observed the same expression of *Pax7* in both satellite cells and type IIB myoneuceli (Supp. Fig. 2).

**Fig. 2.**
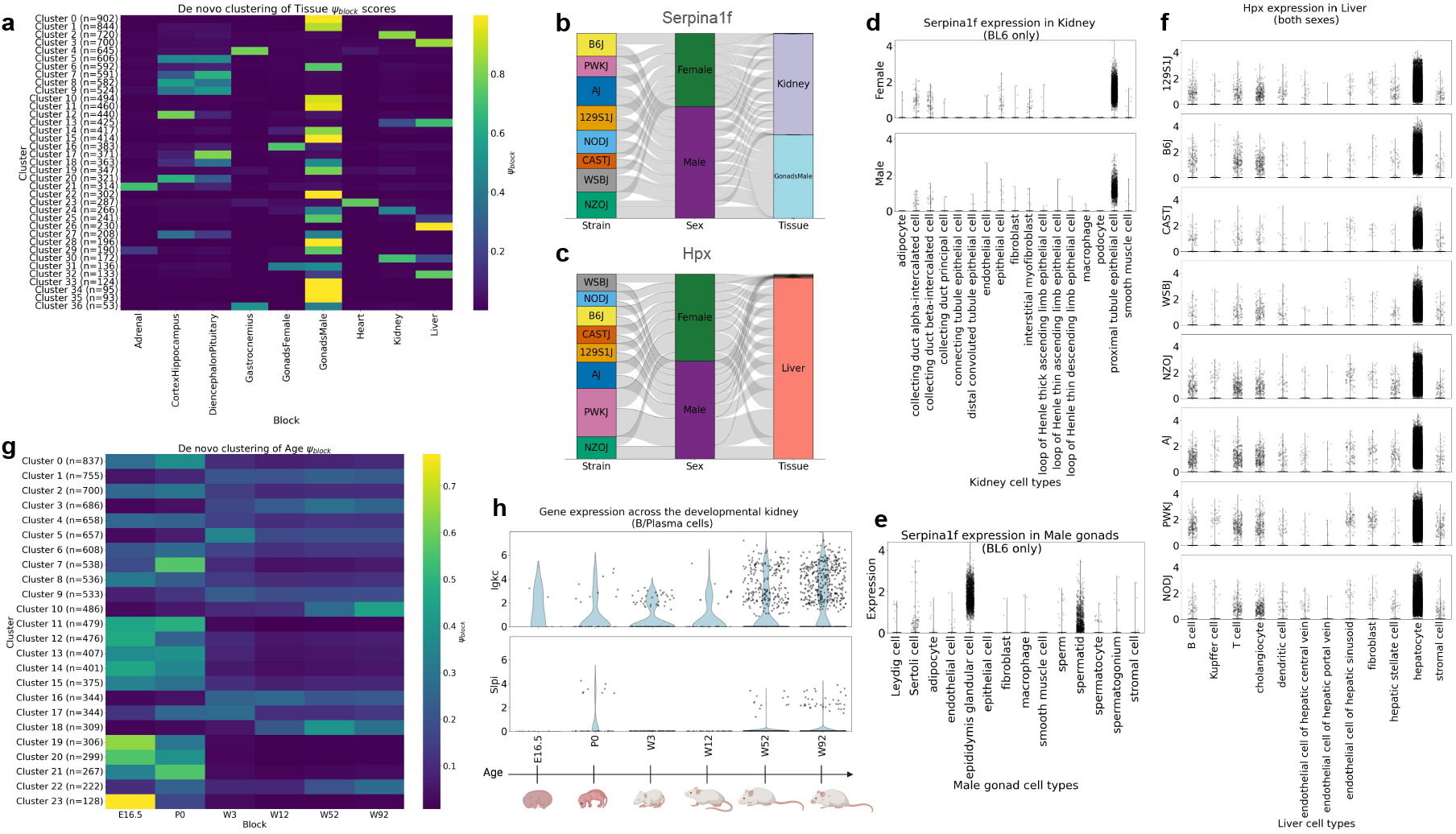
Unsupervised clustering of ψ_block_ scores. a. Heatmap showing de novo unsupervised clustering of *ψ*_*block*_ scores generated from the 8cube dataset (6), partitioned by tissue. **b, c**. Alluvial plots depicting pseudo-bulked gene expression of *Serpina1f* and *Hpx* across strains, sexes and tissues in 8cube (6), generated using wompwomp (29). **d**. Violin plots of *Serpina1f* gene expression in 8cube B6J kidney in each sex, across cell types (6). **e**. Violin plot of *Serpina1f* gene expression in 8cube B6J male gonads across cell types (6). **f**. Violin plots of *Hpx* gene expression in 8cube Liver in both sexes, across strains and cell types (6). **g**. Heatmap showing de novo unsupervised clustering of *ψ*_*block*_ scores generated from the developmental kidney dataset (7), partitioned by age. **h**. Violin plots of *Igkc* and *Slpi* gene expression in the B/Plasma cells of developmental kidney in both sexes, across ages (7), mouse developmental timeline image on x-axis created in https://BioRender.com.

Selecting for genes partitioned by cell type with high Ψ and low *ζ*, we found housekeeping genes for the 8cube gastrocnemius (6) (Fig. 1c). From our analysis, *Akap13*, which encodes an A kinase receptor protein, emerged as one of the top housekeeping genes by cell type within the context of the mouse gastrocnemius (Fig. 1e). We found that *Akap13* outperforms several of the canonical housekeeping genes such as *Gapdh, Actb, Ubc* and *Tbp* as it displays a more uniform distribution across cell types (Fig 1c, 1g). Moreover, it has been documented that *Gapdh* mRNA is highly variable across tissues and cell types (41), making it an unreliable default housekeeping gene across contexts. This result illustrates the advantage of taking a context-aware approach to housekeeping gene selection using ember.

#### Strain-driven cell type specificity

We also uniquely find genes that switch cell type specificity based on strain. *Stk31* encodes serine threonine kinase 31 which enables protein kinase activity. Bao *et al*. 2013 report that *Stk31* is exclusively expressed in spermatogenic cells of testes in mice, their study however is limited to mice of a C57BL/6J background (42). While in the 8cube dataset we observe that *Stk31* expression is generally highest in male gonads, specifically in CASTJ we observe some expression in multiple other tissues (Supp. Fig 1e). For instance in gastrocnemius, we observe that it is selectively expressed in fibro-adipogenic progenitor cells in CASTJ mice but in type IIb myonuclei in PWKJ mice (Fig. 1g). This strain-dependent expression reveals that genetic background can drive cell type specificity, challenging the notion of universal cell type markers. This especially motivates exercising caution when drawing cross-strain and cross-species conclusions from a mouse model. To fuel effective and informed use of mouse models in biological research, we have deposited specificity and gene expression information across the 8 diverse founder strains in 8 tissues into a publicly available and programmatically accessible API database with MCP server integration and a website (*Data Availability*). Together, these results showcase the ability of ember as a tool to evaluate the efficacy of marker and housekeeping genes to a partition while also identifying novel strain-aware cell type-specific genes.

### Unsupervised learning from entropy metrics can identify clusters of specific genes

#### Clustering by tissue

We performed unsupervised machine learning using the leiden clustering algorithm on the entire 8cube data, clustering on *ψ*_*block*_ scores of each tissue (Fig. 2a) (6, 43). We found 4863 out of the 14181 genes with Ψ *>* 0.45, *ζ >* 0.45 were specific to male gonads (34%). This is in agreement with previous work in humans by Djureinovic *et al*. 2014, which confirms that male gonads are the tissue with most tissue-specific genes (44). We also found clusters of genes that are specific to pairs of tissues involving male gonads (Table 1, Supp. tables 2-8). After performing a functional enrichment analysis (45) of the 226 genes with shared specificity to kidney and male gonads, we found 68 genes are predicted to be associated with the *Twist* transcription factor. *Twist1* and *Twist2* are paralogs in the *Twist* subfamily of basic loop-helix-loop proteins, which are essential for embryogenesis and are conserved across vertebrates and invertebrates (46, 47). The role of *Twist1* has been extensively studied in the kidney and is implicated in kidney disease (48–50). However, previous research has found that Twist1 is not detected in the murine testes (51), so these genes with a shared predicted association with the *Twist* family in both male gonads and kidney lead us to a novel line of biological questioning.

**Table 1.** Clusters of genes from Fig 2a. with specificity to male gonads and one other tissue.

| Tissue | Cluster | Gene Count | Functional Enrichment Summary |
| --- | --- | --- | --- |
| Female Gonads | 31 | 136 | mitosis, chromosome segregation, reproductive processes |
| Kidney | 24 | 226 | Twist motif enrichment, peptidase activity, serine-type peptidase activity, serine hydrolase activity |
| Cortex, Hippocampus | 27 | 208 | Cell signaling, membrane receptor activity, GPCR activity, olfactory receptor activity |
| Diencephalon, Pituitary | 18 | 363 | Sensory perception, response to chemical stimulus, GPCR activity, olfactory receptor activity |
| Gastrocnemius | 36 | 53 | None |

*Serpina1f*, which encodes serine peptidase inhibitor, clade A member 1f, is one of the 68 genes specific to male gonads and kidney and predicted to be associated with *Twist* (Fig 2b). *Serpina1f* has been previously identified as kidney-specific in a semi-quantitative PCR analysis study of adipose tissue, muscle, heart, lung, liver and kidney (52). We see from our data that *Serpina1f* is largely localized to proximal tubule epithelial cells in both males and females (Fig 2b, 2d). In male gonads, *Serpina1f* has been localized to mouse epididymis in an RNA in-situ hybridization study (53, 54) and to sperm cell membrane in a proteomics study (55). We also observed localization of *Serpina1f* to epididymis and spermatid in male gonads (Fig 2e). These results validate that *Serpina1f* is reproducibly and distinctly observed in both kidney and male gonads.

The only tissue with a cluster of highly specific genes (*ψ*_*block*_ *>* 0.8) other than male gonads was Liver (Fig 2a, Cluster 26, 230 genes). A functional enrichment analysis on this cluster of genes revealed regulation of large sets of these genes by six transcription factors: *Tcf-1, Cdx-2, Crx, Mef-2c, Sox-10* and *Cdp. Tcf-1* or *Hnf1a* encodes a liver-enriched transcription factor critical to several aspects of liver function (56–60). *Hpx* is one of the 26 genes that we identified as liver-specific and regulated by *Tcf-1* (Fig 2c, Supp. table 3, 8). *Hpx*, encodes hemopexin, a protein that binds heme and transports it to the liver for breakdown and iron recovery. While *Hpx* is visibly specific to liver across all 8 tissues (Fig 2c), at the strain level we noticed that *Hpx* shows a slight bias towards PWKJ mice, which are the most prone to develop hepatic inflammation of the founder strains (61). At the cell type level, we noticed localization of *Hpx* to hepatocytes across all strains (Fig 2f). These results show that we can find clusters of biologically meaningful genes with tissue specificity.

#### Clustering by age

We used ember to understand gene specificity in a developmental kidney dataset produced by Chen *et al*. 2025. In this data, the developing kidney is captured in six stages: E16.5, P0, week 3 (W3), week 12 (W12), week 52 (W52) and week 92 (W92), corresponding approximately to embryo, newborn, youth, adolescence, adult and old age in humans(7). Partitioning this data by age, we generated entropy metrics and then clustered genes by their *ψ*_*block*_ scores (Fig. 2g). These age specificity measurements were calculated across the whole kidney dataset and therefore reflect a combination of age associated changes in cell type composition and within cell type expression. This analysis revealed a cluster of genes specific to old age with high specificity to W92 and slightly lower specificity to W52 (Cluster 10, 486 genes, Supp. table 9). A gene set enrichment analysis using the molecular signatures database (MSigDB) on this cluster of genes revealed heightened immune activity (Supp. table 10) (62). This is in agreement with findings from the Chen *et al*. 2025 authors, who found steadily increasing immune cell proportions from W12 to W92, indicating an accumulation of immune cells in kidneys upon aging (7). One gene in this cluster of *“aging-specific genes”* is *Igkc*, which encodes immunoglobulin kappa constant. *Igkc* has previously been identified as an aging marker across several tissues, including kidney, in a proteomics study comparing 32 and 72 week old mice (63).

We found a cluster of genes specific to non-consecutive developmental stages. Cluster 22, a cluster of 222 genes, shows specificity to ages P0, W52 and W92. A gene set enrichment analysis of this cluster using MsigDB reveled heightened immune and inflammatory response (Supp. table 9, 11) (62). A study by Speer *et al*. 2025 found that neonatal sepsis lead to increased expression of renal tissue inflammation and injury biomarkers, consistent with acute kidney injury, and drawing some parallels between the molecular signatures of a new born mouse to that of an aged one (64), hinting at that biology that may be underlying this unique nonconsecutive bimodal expression. Secretory leukocyte protease inhibitor (*Slpi*), is a gene within this cluster of genes specific to new borns and old age, plays a regenerative role in the ischemic renal (I/R) injury model, through activation induced by *MIF-2/D-DT* (65). Together, these results show that applying unsupervised learning techniques to entropy metrics provides a context-aware methodology for identify clusters of genes with distinct and unexpected expression patterns.

### Tissue and cell type are the dominant source of biological variation in a scRNAseq experiment

#### Tissue-linked specificity

Generating entropy metrics for multiple partitions in a dataset allows unique insights into biological patterns and trends. We generated Ψ and *ζ* values partitioning the entire 8cube data by tissue, sex and mouse strain and their two and three way interaction terms (6). We extracted lists of genes most specific to these partitions (Ψ *>* 0.75, *ζ >* 0.75 to ensure comparable sizes of gene sets) and from these seven sets of genes, we generated an upset plot to visualize gene membership (Fig. 3a) (66). We found genes to be most specific to tissue, along with a large set of genes with sex specificity linked to tissue. We also found that tissue specificity dominates strain specificity, with only 123 genes highly specific to strain and 1402 genes highly specific to tissue. To further validate this observed pattern, we performed a principal component analysis of gene expression data pseudo-bulked by mouse strain and tissue. PC1 and PC2 together account for more than half of the total variability (55.8% of total variance, 34.3% and 21.5% for PC 1 and 2 respectively), indicating that the major biological factors driving variation are captured effectively within the first two principal components (Fig. 3b). The pseudo-bulked expression profiles show distinct clustering by tissue (shape), over mouse strain (color). In addition, a correlation cluster map of this pseudo-bulked data also revealed that tissues across strains had higher correlation to one another (Supp. Fig 3). Together, the principal component analysis and the correlation structure of the pseudo-bulked profiles (Fig. 3b, Supp. Fig. 3) show that tissue is the dominant source of biological variation, making genes with specificity to partitions other than tissue especially interesting.

**Fig. 3.**
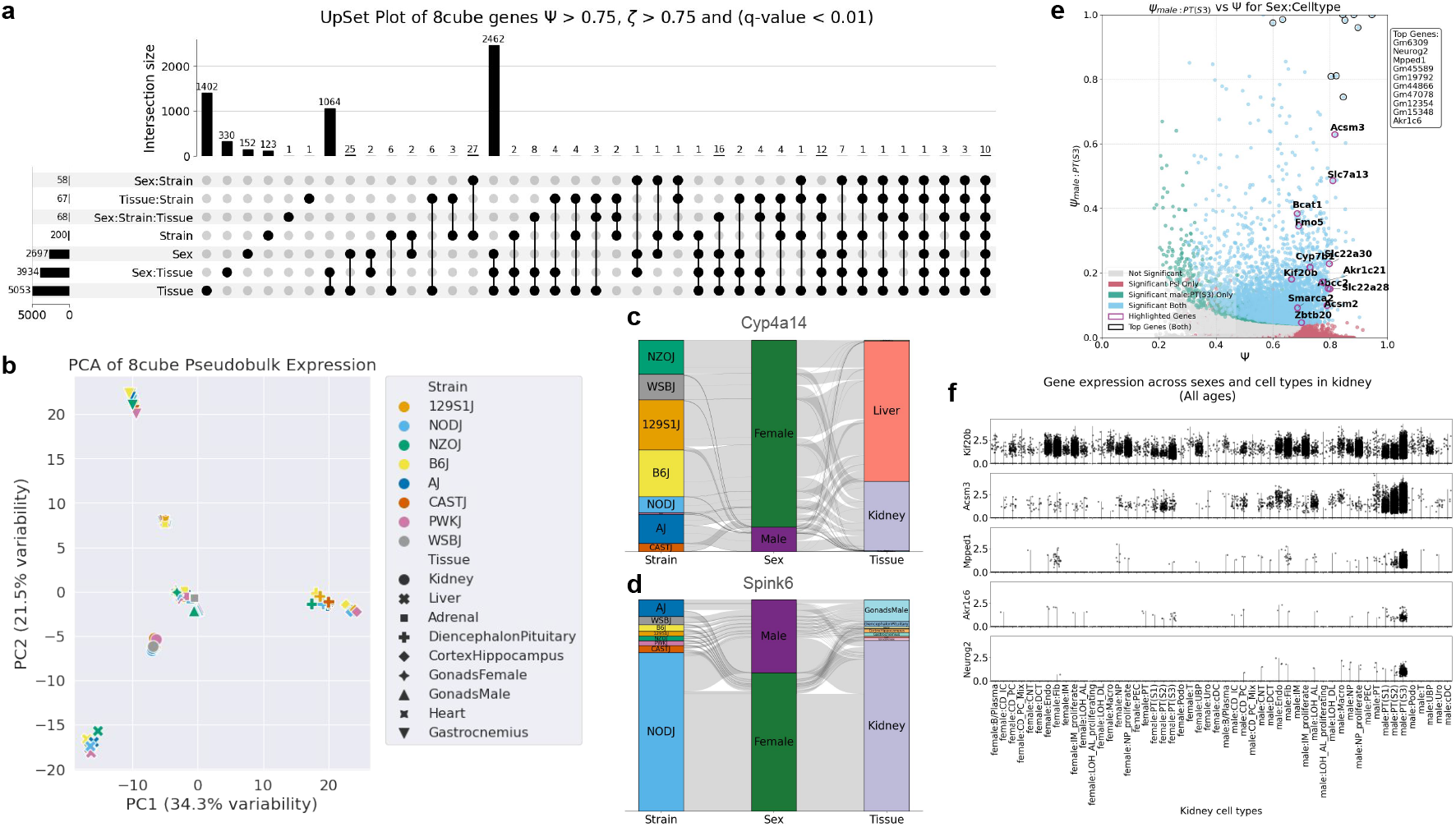
Visualizing specificity patterns across partitions a. Upset plot generated from 8cube dataset (6) selecting highly specific genes partitioned by sex, strain, tissue and their 2-way and 3-way interaction terms. Thresholds used for highly specific gene selection are Ψ *>* 0.75 and *ζ >* 0.75. Global testing correction was performed across all 7 partitioned and genes selected passed a significance threshold of 0.01. **b**. PCA plot of 8cube data pseudo-bulked by strain (color) and tissue (shape) (6). **c, d**. Alluvial plots depicting pseudo-bulked gene expression of *Cyp4a14* and *Spink6* across strains, sexes and tissues in 8cube (6), generated using wompwomp (29). **e**. Scatter plot showing gene specificity to male proximal tubule epithelial cells (Segment 3) in developmental kidney (7), partitioned by sex:celltype. Genes with high Ψ and high *ψ*_*malePT*(*S*3)_ are identified as marker genes (highlighted in black). Top differentially expressed genes in male PT(S3) cells identified by Chen *et al*. 2025 (7) are highlighted in purple. Significance testing via permutation testing for both metrics is shown by color: genes significant for both are light blue and exhibit reliable specificity. Top marker genes are listed in the top-right text box. **f**. Violin plots for *Kif20b* and *Acsm3*, identified as differentially expressed in male PT(S3) cells by (7) and *Mpped1, Akr1c6* and *Neurog2*, identified by ember as highly specific to male PT(S3) cells.

The upset plot in Fig 3a isolates *Cyp4a14*, which encodes a cytochrome P450 monooxygenase, to be the only gene highly specific to sex:strain:tissue over other partitions (Fig. 3c). *Cyp4a14* shows biased expression towards female liver and kidneys in all strains except PWKJ (Fig. 3c). In a bile duct ligation murine model, overexpression of *Cyp4a14* attenuated fibrosis, while knockout aggravated it (67). In this analysis we also found *Spink6* to be the only gene highly specific to tissue:strain over other partitions (Fig 3d). *Spink6* encodes serine peptidase inhibitor kazal type 6 and shows biased expression towards NODJ kidney.

#### Cell-type-linked specificity

We observe a similar pattern at the tissue level. In a principal component analysis of 8cube kidney expression pseudo-bulked by genotype and cell type, PC1 and PC2 together account for 38.8% of total variance (26.2% and 12.6% respectively), and profiles cluster by cell type rather than by strain (Supp. Fig. 4b) (6). In the developing kidney (7), we see that specificity by cell type dominates specificity by age and sex. (Supp. Fig 4a, b, c). In a tissue like the kidney, where it is interesting to study sexual dimorphism, being able to resolve specificity to sex from specificity to sex within a cell type is critical (68). Chen *et al*. 2025 identified that proximal tubule epithelial cells, specifically segment 3 (PTS3) present the largest number of sexually dimorphic genes (7). Chen *et al*. 2025 identified genes as sexually dimorphic in PTS3 cells with biased expression towards males using a non-parametric Wilcoxon rank sum through the Seurat package.

**Fig. 4.**
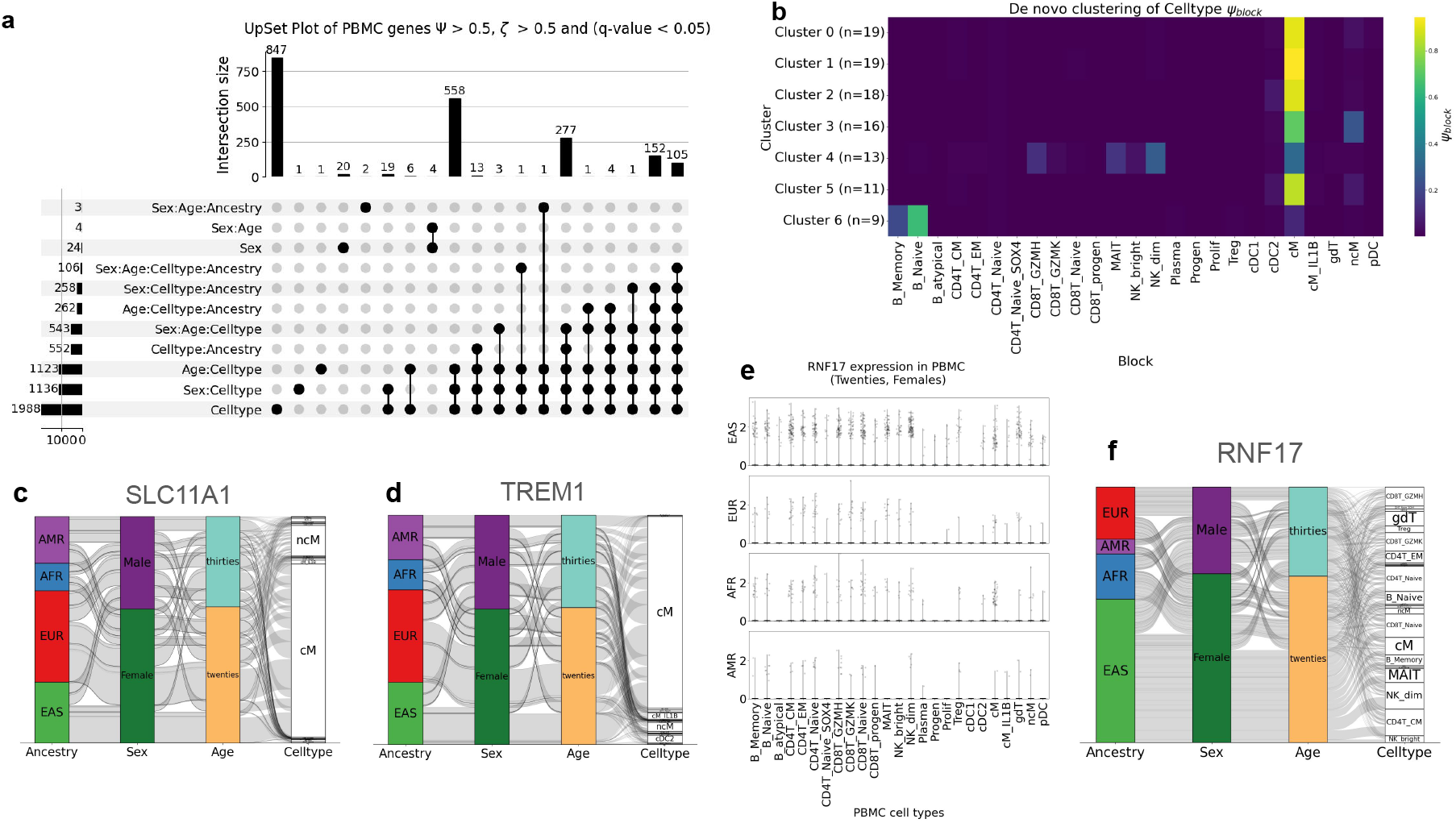
Specificity in human PBMCs collected from 255 individuals from diverse ancestriesa. Upset plot generated from human PBMC dataset selecting highly specific genes partitioned by Sex, Age, Ancestry and Celltype and their 2-way, 3-way and 4-way interaction terms. Thresholds used for highly specific gene selection are Ψ *>* 0.5 and *ζ >* 0.5. Global testing correction was performed across all 15 partitions and genes selected passed a significance threshold of 0.05. **c, d, f**. Alluvial plots depicting pseudo-bulked gene expression of *SLC11A1, TREM1* and *RNF17* respectively across ancestries, sexes, ages and cell types in human PBMCs. **e**. Violin plots of *RNF17* gene expression in 20-29 year-old females across ancestries and cell types in human PBMCs.

We re-analyzed the developmental kidney dataset using ember to understand male-biased sexual dimorphism in PTS3 cells (7). We generated entropy metrics, partitioning the data by sex:celltype and overlayed the top genes identified by Chen *et al*. 2025 on a scatter plot showcasing specificity to male PTS3 cells (Ψ vs. *ψ*_*male*:*PTS*3_). We noticed that these genes had Ψ *>* 0.6, and had a spread of *ψ*_*male*:*PTS*3_ scores lower than 0.65 (Fig. 3e), indicating that while these genes may have high expression in male PTS3s, they are not truly specific to the block. Upon closer inspection we found that one of the genes on this list, *Kif20b* along with having significant expression across various cell types, has biased expression towards males in PTS3 cells but towards females in loop of henle ascending limb proliferating cells, making it specific to neither sex nor cell type (Fig. 3f). We found that another gene on this list *Acsm3*, has high expression not only in male PTS3 cells but also in male PTS1 and male PTS2 (Fig 3f). Conversely, genes with high Ψ and *ψ*_*male*:*PTS*3_ scores identified by ember show distinct and exclusive specificity to male PTS3 cells (Fig 3f). These results re-emphasize the advantage of leveraging decomposable gene-specificity when finding differentially expressed genes in a multivariate dataset.

#### Variation in humans

We generated entropy metrics for a human PBMC dataset of 255 individuals from four different ancestries: European, African, East Asian and Admixed American, partitioning by age, cell type, sex, ancestry and their two-way, three-way and four-way interaction terms (Fig 4a). We found, similar to mouse, that specificity within PBMCs was dominated by cell type and that sex, age and ancestry related specificity are largely linked to cell type. 105 genes were identified as specific to cell type or a cell-type-linked interaction term (Supp. table 12). We clustered these 105 genes on their *ψ*_*block*_ scores and found that 67 of the 105 genes have highest *ψ*_*celltype*_ scores in classical monocytes, suggesting that variation in human PBMCs may be concentrated in this cell type (Fig. 4b). One of the these highly variable genes is *SLC11A1*, which encodes a member of the solute carrier family 11, a family of proton-coupled divalent metal ion transporters (Fig. 4c). Mutations in *SLC11A1* have been associated with susceptibility to diseases such as tuberculosis in Chinese population (69), kawasaki disease in Japanese population (70), cutaneous leishmaniasis in Pakistani population (71), type 2 diabetes mellitus in Iranian population (72) and many other ancestry specific variants that offer resistance or heightened susceptibility to disease (73).

Another gene in this list of 105 highly variable genes is *TREM1*, which encodes triggering receptor expressed on myeloid cells 1 and is an ancestral predictor for malaria susceptibility (Fig. 4d). A study of Columbian population showed lower malaria susceptibility in people of Native American ancestry, slightly higher susceptibility in European ancestry and oscillated between a protective effect and increased risk with no clear prediction in African ancestry (74). Another gene *IL1B* is on their list of ancestral malaria predictor genes as well as on our list of 105 highly variable genes linked to cell type.

Outside of cell-type-linked specificity, we found 20 genes uniquely specific to sex and 4 genes specific to sex:age, of which 18 were Y-chromosome-linked genes and the remaining 6 were X-chromosome-linked. We additionally found one gene, *RNF17*, which showed distinct specificity by sex:age:ancestry (Fig. 4e, f). *RNF17* encodes a protein containing a RING finger domain and has been previously considered testes specific. We found that *RNF17* shows biased expression in East Asian ancestry females in their twenties in human PBMCs. This pattern is supported by the sample-balanced permutation testing described in *Methods*. However, since 380 of the 784 *RNF17*-expressing cells in this group derive from a single donor, we present this finding as a candidate observation.

Together, these results show that an entropy-guided approach is effective in understanding biological variance in human genetic data and can find known and novel partition-specific and highly variable genes. This analysis also draws to attention some limitations of the mouse model in understanding genetic variation in humans (28). We identified strain-specific genes across all tissues as well as within a tissue in the mouse 8cube data (6) (Fig. 1a), but such a parallel does not exist in humans. There are no genes that are exclusively ancestry-specific in human PBMCs. Instead, in humans we observe that ancestry specificity is linked to sex and cell type.

## Discussion

Lists of marker genes or differentially expressed genes are often important takeaways from a scRNAseq experiment, but current methods lack a homogeneous definition of what constitutes a marker gene. An idealistic marker gene for a block in a partition should have distinct and exclusive expression in this block. Yet, even the long-noncoding RNA *Xist/XIST*, which is indispensable for X chromosome inactivation during early development in both humans and mice, has non-zero expression in normal male tissues (75). This brings to question whether such idealism, when trying to understand the complexities of biology, may prevent us from truly learning what the transcriptomic orchestra of scRNAseq has to say.

A similar argument can be made for the definition of a housekeeping gene. An ideal housekeeping gene has been described as stably expressed irrespective of tissue type, developmental stage, cell cycle state, or external signal (76, 77). However, the inherently stochastic and bursty nature of transcription (78, 79) undercuts the existence of such a perfectly invariant housekeeping gene.

Taking a more nuanced and context-aware approach to marker gene and housekeeping discovery, that accounts for the limitations of an experiment, such as tissues sampled, mouse strain used, sexes sampled, human populations sampled, and rarity and resolution of a cell type, may lead to more reproducible biological insights. Specificity simply identifies genes with expression patterns that are particularly concentrated within a biological context, the significance of this pattern can only be learned upon further investigation. Interpretation of candidates identified using ember should be paired with the underlying expression distribution, consistency across biological replicates, prior literature and complementary functional analyses such as gene set enrichment. Our tool ember’s context-aware, entropy-based approach allow for discovery of nuanced marker and housekeeping genes, quantifying specificity without assuming exclusivity.

A further challenge to interpreting nuanced gene expression lies in the availability of appropriate visualization tools. A violin plot allows for visualization at single-cell resolution, but is constrained to display expression within a partition. Alluvial plots allow for effective visualization of a gene’s expression across multiple partitions, though at the cost of single-cell resolution (29, 80). To balance these trade-offs, we use both alluvial and violin plots in tandem. This complementary approach allowed us to highlight genes with intricate specificity uncovered by ember, providing interpretable and biologically meaningful visualizations. More broadly, our work aligns with critiques of visualization in single-cell genomics (81), emphasizing that the interpretability of biological results is inseparable from the limitations and choices of visualization strategy.

Our work provides a methodology to interpret gene specificity that is distribution agnostic, decomposable, robust to replicates and that fully leverages single-cell resolution, but it is not without its limitations. First, ember requires discrete partitioning of cells into blocks and is not equipped to handle continuous variables. We recommend binning continuous variables into biologically meaningful categories to obtain an approximate estimate of related gene specificity. Likewise when interpreting cell type specificity, transitional, ambiguous cell states must be assigned to discrete categories prior to analysis, making the reliability of cell-type annotations an important prerequisite for meaningful biological interpretation. Second, while ember is sensitive to lowly expressed genes, transcripts detected in very few cells yield unstable entropy estimates. As a result, filtering is required to remove genes expressed in fewer than 100 cells, which improves robustness but may inadvertently exclude biologically relevant or rare transcripts. When dealing with imbalanced block sizes such as with rare cell states in a cell-type partitioning, while entropy metrics are robust to uneven block sizes and can identify emergent patterns, obtaining statistically significant estimates becomes difficult (see *Supplementary Note, Uneven cell-size benchmark*). Finally, computation of entropy metrics and p-values through sampling can be time consuming when dealing with dense datasets containing more than one million cells. We recommend using parallel computation and chunking strategies where possible to make analyses more manageable. Collectively, these constraints highlight areas for future methodological improvement.

When quantifying gene specificity in scRNA-seq experiments, a central challenge lies in placing results within a broader biological context. Frameworks that enable generalizable conclusions are essential for ensuring accessibility and reproducibility. Our entropy-based approach is particularly well-suited to revealing such organism-level patterns, distilling high-dimensional single-cell data into interpretable, large-scale conclusions. From our analyses, we found that the dominant source of biological variation within an organism is tissue or cell type. Specifically in the 8cube dataset, we conclude that tissue specificity outweighs genotype specificity. These findings align with conclusions made by Gilad and Mizrahi-Man in their reanalysis of the ENCODE data, that variability of genes by tissue is greater than variability by species (82). Taken together, these results reinforce the view that tissue is a primary driver of transcriptomic specificity.

Despite this broader pattern of specificity, it is wise to exercise caution when drawing cross-strain and cross-species conclusions. We observe in our analysis of the 8cube data (6) that there are many distinct differences in gene expression between strains. Strain-specific diversity limits the extent to which findings can be generalized. The heavy reliance on inbred strains exacerbates this issue, raising the possibility that observed specificity reflects artifacts of restricted genetic backgrounds rather than broadly conserved mechanisms. These concerns emphasize the danger of extrapolating from a single strain to complex, heterogeneous populations. We have deposited specificity and gene expression information across the 8 diverse founder strains into a publicly available and programmatically accessible API database with MCP server integration and website (*Data Availability*), fueling the effective and informed use of mouse models in biological research.

An important evolutionary question emerges from these observations: how do highly specific genes arise and change across lineages? Is there an adaptive advantage to maintaining a housekeeping role, or can specificity confer selective benefits? Moreover, how can a gene that is highly specialized in one mouse strain acquire a different function and specificity in another? The evolutionary trajectories of marker and housekeeping genes remain poorly understood (14, 83, 84). Addressing these questions may provide key insights into the origins of specificity and the selective pressures shaping gene expression programs across species. By providing a robust and quantitative framework for measuring and dissecting gene specificity, ember offers a way to begin tracing such evolutionary trajectories, enabling comparative analyses of how marker and housekeeping functions emerge, shift, and are maintained across contexts.

## Conclusions

We introduce ember, an entropy-based framework for quantifying gene specificity in multivariate single-cell data that is decomposable and distribution-agnostic. By grounding specificity in information theory, ember enables interpretable analysis of gene expression across complex partitions.

Applying ember to mouse and human datasets, we show that gene specificity is highly context-dependent and frequently defies canonical expectations. We uncover strain-dependent switching of cell type specificity, shared specialization across tissues, and multivariate specificity across developmental time and ancestry. At a systems level, tissue and cell type emerge as the dominant drivers of transcriptomic variation, while genetic background and demographic variables introduce secondary, context-dependent effects. To support further exploration, we provide a comprehensive mouse specificity database with an accompanying web interface, API, and MCP server.

More broadly, our entropy-based framework reframes gene specificity as a continuous, context-aware property rather than a fixed attribute. This shift enables systematic and scalable interrogation of gene regulation across diverse biological settings and clarifies the limitations of interpreting markers and model systems in isolation. By unifying theoretical rigor with practical applicability, ember establishes a generalizable foundation for studying cellular identity and the impact of genetic variation across organisms.

## Methods

**Ψ, *ζ*, and *ψ***_**block**_ **scores**. Let *x*_*i*_ be the normalized count of a gene in cell *i*. Define the probability

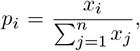

i.e., the proportion of the gene’s total count found in cell *i*.

The **total entropy *E***_*T*_ of the gene expression across all cells is defined as:

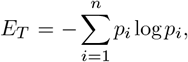

where *n* is the number of cells.

Let *n* cells be partitioned into *r* blocks. We define Ψ based on the within-block and between-block decomposition of entropy. And let *q*_*jC*_ be the proportion of expression in cell *j* relative to the total in block *C*:

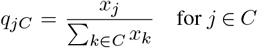

Then, ***E***_*C*_ is the **entropy within the block *C***:

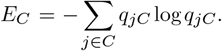

The total **within entropy *E***_*W*_ is the weighted sum, where *p*_*C*_ is the proportion of expression across all cells in block *C*:

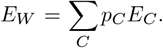

Since the total entropy *E*_*T*_ can be decomposed as:

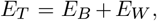

Where **between entropy *E***_*B*_ of the gene expression pseudobulked by block is defined as:

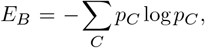

Then, we define **Ψ (Psi) as the fraction of gene-expression entropy retained within partition blocks** of *n* cells:

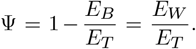

For *E*_*T*_ *>* 0, Ψ = 1 when all gene expression is contained in a single block. Conversely, Ψ = 0 when no entropy remains within blocks, i.e., expression in every occupied block is concentrated in a single cell.

We define the normalized contribution of each block to *E*_*W*_ as ***ψ***_*block*_, a measure of **specificity to a block**. Then *ψ*_*block*_ for block *C* is given by:

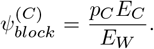

Because *E*_*W*_ = *C p*_*C*_*E*_*C*_, *ψ*_*block*_ scores are non-negative and sum to one across blocks, providing a compositional quantification of where gene expression is concentrated across a partition.

Let *ψ*_blocks_ denote the set of *ψ*_*block*_ scores across all (r) blocks:

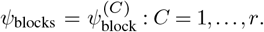

We define ***ζ*** as the normalized Kullback-Leibler divergence of the distribution of *ψ*_*blocks*_ to the uniform distribution. This metric quantifies **specificity to a partition**.

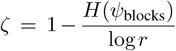

*ζ* = 0 represents perfect alignment with a uniform distribution of *ψ*_block_ scores in a partition and *ζ* = 1 indicates perfect specificity to a single block in a partition.

### Sampling over replicates

To ensure balanced representation of replicates across categories in an experimental design, we implemented a sampling strategy that generates subsets of samples with equal representation of experimental conditions within each category. For every unique category (e.g., genotype, strain, or age group), replicates were iteratively selected so that each condition (e.g., sex) was represented exactly once per sampling draw. The distinction between “category” and “condition” was made for conceptual clarity, although these variables can be defined interchangeably. Sampling was performed at the level of unique sample identifiers recorded in the sample metadata (e.g., mouse_ids or sample_-ids). Multiple balanced subsets (100 draws except for the across tissue analysis where 50 draws were used to speed up calculations) were generated while minimizing repeated selection of the same samples across draws by tracking sample usage frequency. Random seeds were used to ensure reproducibility, and the approach enforced complete balance by requiring at least one sample for every category-condition pair. (Supp. Fig 5 a,b,c)

Arithmetic mean and standard deviation values for Ψ and *ζ* were calculated to capture variation in specificity across biological replicates.

Since *ψ*_block_ values represent compositional fractions that sum to one within each partition, we aggregated replicate level *ψ*_block_ estimates across sampling draws using compositional data analysis principles. Specifically, the Aitchison mean was used to compute the central tendency of *ψ*_block_ values across draws, while the geometric standard deviation quantified variability in block-specific specificity estimates.

### Permutation testing to generate p-values and q-values

Significance of Ψ, *ψ*_block_, and *ζ* values were estimated by comparing observed values to null distributions generated from permutation testing. For each of *n*_iterations_ permutations, a balanced subset containing exactly one biological replicate per category–condition combination was drawn using the same replicate-balancing procedure described above (*Sampling over replicates*). Partition labels were then permuted among the cells within this balanced subset, and the entropy metrics were recomputed on the permuted data. Because at most one replicate represents any category within a given draw, and because the observed statistics are computed under the same balanced-sampling scheme, no single replicate can dominate a permutation by virtue of its cell count. Label permutation itself is performed at the cell level within each balanced draw rather than by reassigning replicate identity. We used *n*_iterations_ = 1,000 permutations throughout. For each gene, empirical *p*-values were computed as 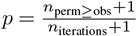, giving a minimum attainable *p*-value of 1*/*1001 ≈ 0.000999.

All resulting *p*-values were combined across metrics and subjected to global false discovery rate (FDR) correction using the Benjamini–Hochberg procedure to obtain *q*-values. Correction was performed in two stages: within a single partition, Ψ and *ζ* (and *ψ*_block_, where computed) *p*-values were pooled and corrected together; for the multi-partition summary analyses, *p*-values were pooled a second time across all partitions analyzed in that figure prior to correction (7 partitions for Fig. 3; 15 partitions — Sex, Age, Ancestry, Cell type and their 2-, 3- and 4-way combinations — for Fig. 4).

### Octagon plot representation of block specificity

The octagon plot from Fig. 1a was designed to visualize variation in specificity across biological replicates to the 8 different founder mouse strains. Because *ψ*_block_ values for a given partition represent compositional fractions that sum to one, gene-specificity profiles can be represented as points within a simplex. In this framework, each coordinate corresponds to the proportion of a gene’s specificity attributed to a particular block, such that all coordinates are non-negative and collectively sum to unity. In general, an *n*-simplex can be projected onto a regular (*n* + 1)-gon, yielding *n*!*/*2 unique 2D projections. For visualization of strain-level specificity across the eight founder mouse strains, the geometry of the 7-simplex was projected into two dimensions as one of 2,520 possible regular octagonal representations. We chose the projection ordered by distance in single nucleotide variants to B6J mice from Ferraj *et al. 2023* (27). It should be noted that distances in this projection are not necessarily exact distances in the original compositional space.

The genes displayed were drawn from those identified as strain-specific in the strain partition using a relaxed cutoff of 0.45 rather than the default 0.5. This value was chosen on the basis of prior knowledge of an established strain-specific gene, *Cwc22* (30) which has *ζ* = 0.46 and Ψ = 0.90 in this partition (both *q − value <* 0.003) and would have been excluded at the default cutoff, as would *Or5p58* (*ζ* = 0.47) and *Or13p8* (*ζ* = 0.49). Data used to generate this plot is available at the supplementary data link (*Supplementary Information*). Code to generate this compositional octagon plot is in the *Code Availability* section.

### Data normalization

All entropy metrics were computed on depth-normalized, log-transformed expression. ember applies no normalization internally. Human PBMC counts were normalized to 10,000 counts per cell and natural-log transformed using `scanpy.pp.normalize_total` and `scanpy.pp.log1p` (85), with raw counts retained in `layers[‘counts’]`. The 8cube data were quantified with kallisto/bustools, corrected for ambient RNA with Cell-Bender, and normalized by total UMI count followed by logarithmic transformation in Scanpy by the original data generators (6, 85). The developmental kidney data were aligned with Cell Ranger ARC (mm10) and log-normalized in Seurat by the original data generators (7, 86).

### Pre-processing: Gene and block filtering

For every dataset, genes detected in fewer than 100 cells were removed prior to entropy computation. In the human PBMC data this excluded 9,689 of the 36,601 genes in the GRCh38/GENCODE 32 reference (26.5%), retaining 26,912. In the 8cube gastrocnemius data it excluded 12,397 of 37,427 genes (33.1%), retaining 25,030. The corresponding retained-gene counts were 27,903 for 8cube kidney, 41,786 for 8cube across all tissues, and 20,069 for the developmental kidney data. No minimum block size was applied: all observed blocks contribute to the entropy metrics irrespective of cell number, and the only structural requirement is that each block contain at least one biological replicate, enforced by the balanced replicate sampler. In the human PBMC data all 400 blocks of the full Sex *×* Age *×* Cell type *×* Ancestry partition were populated, as were all lower-order partitions. In the 8cube data, 2 of 18 Sex *×* Tissue and 16 of 144 Sex *×* Strain *×* Tissue combinations are absent because the gonad tissues are sex-exclusive; these are biologically impossible combinations rather than filtered blocks. All 25 manually annotated PBMC cell types were retained, including the rarest: cDC1; 937 cells, 0.12% of cells, detected in 232 of 255 donors.

### Interpreting entropy metrics

a. **Specific to a block / “Marker” genes** Metrics *ψ*_block_ and Ψ were used to identify genes with high specificity to individual blocks. Genes with the highest *ψ*_block_ and Ψ values were considered most specific to the selected block (e.g., skeletal muscle satellite cells in gastrocnemius, Fig. 1b).
b. **Specific to a partition** Metrics *ζ* and Ψ were used to identify genes specific to a partition. Genes with the highest *ζ* and Ψ values were interpreted as most specific to the partition (e.g., gastrocnemius partitioned by cell type, Fig. 1c).
c. **Non-specific to a partition / “Housekeeping” genes** Metrics *ζ* and Ψ were also used to identify non-specific or “housekeeping” genes. Genes with low *ζ* but high Ψ values were interpreted as least specific to any single partition (e.g., gastrocnemius partitioned by cell type, Fig. 1c).

### Choosing Ψ and ζ threshold

Entropy-based specificity metrics were calculated using the ember Python package (see *Code Availability* for repository link). Gene-level entropy metrics were first computed for each dataset, after which genes with *q − values <* 0.05 were retained. Genes were then ranked according to their respective entropy metrics, and biologically meaningful thresholds (default 0.5 if no prior knowledge) were applied to identify the most specific genes within each partition or block. Because specificity is relative to the biological partition and dataset being analyzed, these thresholds are intended as context-dependent filtering criteria rather than universal cutoffs. Thresholds may be relaxed to retain biologically relevant genes that fall near the default cutoff or made more stringent when restricting an analysis to the most highly specific genes or reducing the size of large gene sets for downstream analyses.

Both cases arise in this work. We relaxed the cutoff to 0.45 for the strain-specific gene selection underlying Fig. 1a, for the reasons given in that section. We retained the same value for the tissue clustering analysis in Fig. 2a. Since the clustering separates genes by their *ψ*_block_ profiles, restricting the input to only the most extreme values is not desirable. Conversely, we made the cutoff more stringent (0.75, with *q − value <* 0.01) for the analysis across all eight tissues, where the number of highly specific genes in each partition was too large to inspect directly. At 0.75 the selection is restricted to genes that are very highly specific to a single block within a partition, reducing the set from 13,773 to 5,699 genes.

### Finding specificity trends using upset plots

To identify large-scale trends in gene specificity across multiple partitions, we computed Ψ and *ζ* scores for all relevant partitions, including interaction terms (e.g., two-way, three-way, or four-way combinations). Genes with *q − value <* 0.05 for both metrics were selected. Genes exhibiting strong specificity within each partition were selected using a biologically informed or default cutoff of 0.5 when no prior information was available. Accordingly, the human PBMC upset analysis used the default cutoff of 0.5, whereas the 8cube analysis across tissues used the more stringent 0.75 with *q − value <* 0.01 to keep the number of intersecting sets interpretable across seven partitions. The sets of highly specific genes from each partition were then combined to define gene memberships, which were visualized using upset plots to reveal overlaps and sets of genes with shared specificity patterns across partitions (66). Data used to generate upset plots is available at the supplementary data link (*Supplementary Information*). Code to generate upset plots is in the *Code Availability* section.

## Declarations

## Abbreviations

scRNA-seq: single cell RNA sequencing
snRNA-seq: single nucleus RNA sequencing
TSI: Tissue Specificity index
PEM: preferrential expression measure
WSBJ: WSBEiJ
NZOJ: NZO/HlLtJ
CASTJ: CASTEiJ
B6J: C57BL/6J
129S1J: 129S1/SvImJ
PWKJ: PWK/PhJ
NODJ: NOD/ShiLtJ
PTS3: Proximal tubule epithelial cells segment 3

## Ethics approval and consent to participate

Not applicable

## Consent for publication

Not applicable

## Availability of Data and Materials

Mouse specificity explorer website for visualizing specificity and gene expression in the 8cube data (6): https://mouseexplorer.onrender.com. 8cubeDB API documentation: https://eightcubedb.onrender.com/docs. 8cubeDB API, MCP server and mouse explorer website source code: https://github.com/pachterlab/8cubeDB. ember python package and documentation: https://github.com/pachterlab/ember.

## Data Availability

The accessibility details of the 8cube dataset are described in the Data Availability section of (6) and of the developmental kidney dataset in the Data Availability section of (7). The Ham *et al*. 2025 data accessibility details are described in their Data Availability section (40).

The human PBMC data were generated by the Ye lab (UCSF) as part of IGVF (award HG012076) and are available from the IGVF portal: 75 multiplexed 10x Multiome wells (`XHLT1-POOL-XX{n}-SC{m}`) spanning 335 donors, accessioned per well as `IGVFDS` file sets (measurement sets holding raw reads, analysis sets holding the gene-by-cell count matrices, GRCh38/GENCODE 32, released 2026-08-19). Donor records with sex and self-reported ethnicity are `IGVFDO` accessions and the per-donor, per-well biosamples carrying age are `IGVFSM` accessions (all released 2025-09-22). Count matrices and barcode-to-sample maps are openly downloadable; raw reads and ATAC fragments are controlled-access under institutional certificate IP115-26 (data-use limitation GRU). We used the RNA modality only, restricted to donors aged 20–39, the two bins with the most biological replicates (156 and 101 of 335 donors; all other decades ≤ 44), giving 255 donors and 817,684 cells. Per-cell sex, age, ancestry and cell-type annotations are in `Human_-PBMC/metadata.h5ad` in the Zenodo deposit.

## Code availability

The code for reproducing the figures and analyses including benchmark experiment in the *Supplemental Note*: https://github.com/pachterlab/SBRGKLAYWMP_2025. Data to reproduce the figures and analyses: https://doi.org/10.5281/zenodo.17654220.

## Author contributions

N.P.S. contributed to conceptualization, performed analyses, developed the ember python package, mouse explorer website, 8cubeDB API and MCP server, made figures, and drafted the manuscript. S.B. contributed to conceptualization. E.R. generated the 8-cube founder dataset and assisted with cell type annotations. M.G.G., P.K., T.L, M.A., and C.J.Y. generated the human PBMC dataset and performed cell type annotations. B.W. and A.M. contributed to supervision, data acquisition, and funding acquisition. L.P. contributed to conceptualization, supervision, and funding acquisition. All authors contributed to manuscript review.

## Acknowledgments

We thank Conrad Oakes and Joseph Rich for help with optimizing alluvial plots. We thank Ruth Pachter and Juliet Lee for help with interpreting the coding and protein sequences for *Tas1r1*.

## References

1. Junyue Cao, Jonathan S. Packer, Vijay Ramani, Darren A. Cusanovich, Chau Huynh, Riza Daza, Xiaojie Qiu, Choli Lee, Scott N. Furlan, Frank J. Steemers, Andrew Adey, Robert H. Waterston, Cole Trapnell, and Jay Shendure. Comprehensive single-cell transcriptional profiling of a multicellular organism. Science, 357(6352):661–667, 2017. doi: 10.1126/science.aam8940.

2. Alexander B. Rosenberg, Charles M. Roco, Richard A. Muscat, Anna Kuchina, Paul Sample, Zizhen Yao, Lucas T. Graybuck, David J. Peeler, Sumit Mukherjee, Wei Chen, Stephanie H. Pun, David L. Sellers, Bosiljka Tasic, and Georg Seelig. Single-cell profiling of the developing mouse brain and spinal cord with split-pool barcoding. Science, 360(6385):176–182, 2018. doi: 10.1126/science.aam8999.

3. Shristi Pandey, Kimberle Shen, Seung-Hye Lee, Christopher J. Bohlen, Tracy J. Yuen, Brad A. Friedman, et al. Disease-associated oligodendrocyte responses across neurode-generative diseases. Cell Reports, 40(8):111234, 2022. doi: 10.1016/j.celrep.2022.111234. Available at http://research-pub.gene.com/OligoLandscape/.

4. Stefan Salcher, Gregor Sturm, Lena Horvath, Dominik Wolf, Andreas Pircher, and Zlatko Trajanoski. High-resolution single-cell atlas reveals diversity and plasticity of tissue-resident neutrophils in non-small cell lung cancer. Cancer Cell, 40(12):1503–1520.e8, 2022. doi: 10.1016/j.ccell.2022.10.010.

5. Austin D. Reed, Sara Pensa, Adi Steif, Jack Stenning, Daniel J. Kunz, Linsey J. Porter, Kui Hua, Peng He, Alecia-Jane Twigger, Abigail J. Q. Siu, Katarzyna Kania, Rachel Barrow-McGee, Iain Goulding, Jennifer J. Gomm, Valerie Speirs, J. Louise Jones, John C. Marioni, and Walid T. Khaled. A single-cell atlas enables mapping of homeostatic cellular shifts in the adult human breast. Nature Genetics, 56:652–662, 2024. doi: 10.1038/s41588-024-01726-x. Open access.

6. Elisabeth Rebboah, Ryan Weber, Elnaz Abdollahzadeh, Nikhila Swarna, Delaney K. Sullivan, Diane Trout, Fairlie Reese, Heidi Yahan Liang, Ghassan Filimban, Parvin Mahdipoor, Margaret Duffield, Romina Mojaverzargar, Erisa Taghizadeh, Negar Fattahi, Negar Mojgani, Haoran Zhang, Rebekah K. Loving, Maria Carilli, A. Sina Booeshaghi, Shimako Kawauchi, Ingileif B. Hallgrímsdóttir, Brian A. Williams, Grant R. MacGregor, Lior Pachter, Barbara J. Wold, and Ali Mortazavi. Systematic cell-type resolved transcriptomes of 8 tissues in 8 lab and wild-derived mouse strains captures global and local expression variation. Cell Genomics, 6(4), 2026. doi: 10.1016/j.xgen.2026.100964.

7. Siqi Chen, Ruiyang Liu, Chia-Kuei Mo, Michael C. Wendl, Andrew Houston, Preet Lal, Yanyan Zhao, Wagma Caravan, Andrew T. Shinkle, Atieh Abedin-Do, Nataly Naser Al Deen, Kazuhito Sato, Xiang Li, André Luiz N. Targino da Costa, Yize Li, Alla Karpova, John M. Herndon, Maxim N. Artyomov, Joshua B. Rubin, Sanjay Jain, Xue Li, Sheila A. Stewart, Li Ding, and Feng Chen. Multi-omic and spatial analysis of mouse kidneys highlights sex-specific differences in gene regulation across the lifespan. Nature Genetics, 57:1213–1227, 04 2025. doi: 10.1038/s41588-025-02161-x.

8. Ailsa M. Jeffries, Tianxiong Yu, Jennifer S. Ziegenfuss, Allie K. Tolles, Christina E. Baer, Cesar Bautista Sotelo, Yerin Kim, Zhiping Weng, and Michael A. Lodato. Single-cell transcriptomic and genomic changes in the ageing human brain. Nature, 2025. doi: 10.1038/s41586-025-09435-8.

9. David Lähnemann, Johannes Köster, Ewa Szczurek, Davis J. McCarthy, Stephanie C. Hicks, Mark D. Robinson, Catalina A. Vallejos, Kieran R. Campbell, Niko Beerenwinkel, Ahmed Mahfouz, Luca Pinello, Pavel Skums, Alexandros Stamatakis, Camille Stephan-Otto Attolini, Samuel Aparicio, Jasmijn Baaijens, Marleen Balvert, Buys de Barbanson, Antonio Cappuccio, Giacomo Corleone, Bas E. Dutilh, Maria Florescu, Victor Guryev, Rens Holmer, and Alexander Schönhuth. Eleven grand challenges in single-cell data science. Genome Biology, 21:31, 2020. doi: 10.1186/s13059-020-1926-6.

10. Malte D. Luecken and Fabian J. Theis. Current best practices in single-cell rna-seq analysis: a tutorial. Molecular Systems Biology, 15(6):e8746, 2019. doi: 10.15252/msb.20188746.

11. Tim Stuart and Rahul Satija. Integrative single-cell analysis. Nature Reviews Genetics, 20: 257–272, 2019. doi: 10.1038/s41576-019-0093-7.

12. Blanca Pijuan-Sala, Carolina Guibentif, and Berthold Göttgens. Single-cell transcriptional profiling: a window into embryonic cell-type specification. Nature Reviews Molecular Cell Biology, 19(6):399–412, 2018. doi: 10.1038/s41580-018-0002-5.

13. Zoe A. Clarke, Tallulah S. Andrews, Jawairia Atif, Delaram Pouyabahar, Brendan T. Innes, Sonya A. MacParland, and Gary D. Bader. Tutorial: guidelines for annotating single-cell transcriptomic maps using automated and manual methods. Nature Protocols, 16(6):2749–2764, 2021. doi: 10.1038/s41596-021-00534-0.

14. Itai Yanai, Hila Benjamin, Michael Shmoish, Vered Chalifa-Caspi, Maxim Shklar, Ron Ophir, Arren Bar-Even, Shirley Horn-Saban, Marilyn Safran, Eytan Domany, Doron Lancet, and Orit Shmueli. Genome-wide midrange transcription profiles reveal expression level relationships in human tissue specification. Bioinformatics, 21(5):650–659, 2005. doi: 10.1093/bioinformatics/bti042. All data available at http://genecards.weizmann.ac.il/genenote/; GEO accession GSE803.

15. Anupama Roy, Himanshushekhar Chaurasia, Baibhav Kumar, Naina Kumari, Sarika Jaiswal, Manish Srivastava, Mir Asif Iquebal, Ulavappa B. Angadi, and Dinesh Kumar. Featl: a comprehensive web-based expression atlas for functional genomics in tropical and subtropical fruit crops. BMC Plant Biology, 24:890, 2024. doi: 10.1186/s12870-024-05022-9.

16. Lukasz Huminiecki, Andrew T. Lloyd, and Kenneth H. Wolfe. Congruence of tissue expression profiles from gene expression atlas, sagemap and tissueinfo databases. BMC Genomics, 4:31, 2003. doi: 10.1186/1471-2164-4-31.

17. Alec Barrett, Erdem Varol, Alexis Weinreb, Seth R. Taylor, Rebecca M. McWhirter, Cyril Cros, Berta Vidal, Manasa Basaravaju, Abigail Poff, Marc Hammarlund, et al. Integrating bulk and single cell rna-seq refines transcriptomic profiles of individual C. elegans neurons. eLife, v1, 2025. doi: 10.7554/eLife.106183.1. Reviewed Preprint, published April 11, 2025.

18. Corrado Gini. Variabilità e mutabilità: Contributo allo studio delle distribuzioni e delle relazioni statistiche. Tipografia di Paolo Cuppini, Bologna, Italy, 1912.

19. Lidia Ceriani and Paolo Verme. The origins of the gini index: extracts from variability and mutability (1912) by corrado gini. The Journal of Economic Inequality, 10:421–443, 2012. doi: 10.1007/s10888-011-9188-x.

20. Steve O’Hagan, Marina Wright Muelas, Philip J. Day, Emma Lundberg, and Douglas B. Kell. Genegini: Assessment via the gini coefficient of reference “housekeeping” genes and diverse human transporter expression profiles. Cell Systems, 6(2):230–244.e1, 2018. doi: 10.1016/j.cels.2018.01.002.

21. Nadezda Kryuchkova-Mostacci and Marc Robinson-Rechavi. A benchmark of gene expression tissue-specificity metrics. Briefings in Bioinformatics, 18(2):205–214, 2017. doi: 10.1093/bib/bbw008.

22. Michael I. Love, Wolfgang Huber, and Simon Anders. Moderated estimation of fold change and dispersion for rna-seq data with deseq2. Genome Biology, 15:550, 2014. doi: 10.1186/s13059-014-0550-8.

23. Gabriel E. Hoffman and Eric E. Schadt. variancepartition: interpreting drivers of variation in complex gene expression studies. BMC Bioinformatics, 17(1):483, 2016. doi: 10.1186/s12859-016-1323-z.

24. Dmitrii Konstantinovich Faddeev. On the concept of entropy of a finite probabilistic scheme. Uspekhi Matematicheskikh Nauk, 11(1):227–231, 1956.

25. A. Ya. Khinchin. Mathematical Foundations of Information Theory. Dover Publications, Mineola, New York, 2013. ISBN 9780486318448.

26. Michael J. Casey, Jörg Fliege, Rubén J. Sánchez-García, and Ben D. MacArthur. An information-theoretic approach to single cell sequencing analysis. BMC Bioinformatics, 24: 311, 2023. doi: 10.1186/s12859-023-05424-8.

27. Ardian Ferraj, Peter A. Audano, Parithi Balachandran, Anne Czechanski, Jacob I. Flores, Alexander A. Radecki, Varun Mosur, David S. Gordon, Isha A. Walawalkar, Evan E. Eichler, Laura G. Reinholdt, and Christine R. Beck. Resolution of structural variation in diverse mouse genomes reveals chromatin remodeling due to transposable elements. Cell Genomics, 3(5):100291, 2023. doi: 10.1016/j.xgen.2023.100291.

28. Michael C. Saul, Vivek M. Philip, Laura G. Reinholdt, Elissa J. Chesler, and Center for Systems Neurogenetics of Addiction. High-diversity mouse populations for complex traits. Trends in Genetics, 35(7):501–514, July 2019. doi: 10.1016/j.tig.2019.04.003.

29. Joseph Rich, Conrad Oakes, and Lior Pachter. Optimizing alluvial plots. arXiv preprint arXiv:2509.03761, September 2025. doi: 10.48550/arXiv.2509.03761.

30. John P. Didion, Andrew P. Morgan, Amelia M.-F. Clayshulte, Rachel C. McMullan, Liran Yadgary, Petko M. Petkov, Timothy A. Bell, Daniel M. Gatti, James J. Crowley, Kunjie Hua, David L. Aylor, Ling Bai, Mark Calaway, Elissa J. Chesler, John E. French, Thomas R. Geiger, Terry J. Gooch, Theodore Jr. Garland, Alison H. Harrill, Kent Hunter, Leonard McMillan, Matt Holt, Darla R. Miller, Deborah A. O’Brien, Kenneth Paigen, Wenqi Pan, Lucy B. Rowe, Ginger D. Shaw, Petr Simecek, Patrick F. Sullivan, Karen L. Svenson, George M. Weinstock, David W. Threadgill, Daniel Pomp, Gary A. Churchill, and Fernando Pardo-Manuel de Villena. A multi-megabase copy number gain causes maternal transmission ratio distortion on mouse chromosome 2. PLoS Genetics, 11(2):e1004850, 02 2015. doi: 10.1371/journal.pgen.1004850.

31. Anneke Gässler, Charline Quiclet, Oliver Kluth, Pascal Gottmann, Kristin Schwerbel, Anett Helms, Mandy Stadion, Ilka Wilhelmi, Wenke Jonas, Meriem Ouni, Frank Mayer, Joachim Spranger, Annette Schürmann, and Heike Vogel. Overexpression of gjb4 impairs cell proliferation and insulin secretion in primary islet cells. Molecular Metabolism, 41:101042, 06 2020. doi: 10.1016/j.molmet.2020.101042.

32. Yoko Kusuhara, Ryusuke Yoshida, Tadahiro Ohkuri, Keiko Yasumatsu, Anja Voigt, Sandra Hübner, Katsumasa Maeda, Ulrich Boehm, Wolfgang Meyerhof, and Yuzo Ninomiya. Taste responses in mice lacking taste receptor subunit t1r1. The Journal of Physiology, 591(7): 1967–1985, 2013. doi: 10.1113/jphysiol.2012.236604.

33. Anthony Sclafani, Austin S. Vural, and Karen Ackroff. Cast/eij and c57bl/6j mice differ in their oral and postoral attraction to glucose and fructose. Chemical Senses, 42(3):259–267, 02 2017. doi: 10.1093/chemse/bjx003.

34. Junli Liu, Qilin Li, Yixuan Hu, Yi Yu, Kai Zheng, Dengfeng Li, Lexin Qin, and Xiaochun Yu. The complete telomere-to-telomere sequence of a mouse genome. Science, 386(6726): 1141–1146, 12 2024. doi: 10.1126/science.adq8191.

35. Bailey Francis, Landen Gozashti, Kevin Costello, Takaoki Kasahara, Olivia S. Harringmeyer, Jingtao Lilue, Mohab Helmy, Tadafumi Kato, Anne Czechanski, Michael A. Quail, Iraad Bonner, Emma Dawson, Anne Ferguson-Smith, Laura Reinholdt, David J. Adams, and Thomas M. Keane. The structural diversity of telomeres and centromeres across mouse subspecies revealed by complete assemblies. bioRxiv, 10 2024. doi: 10.1101/2024.10.24.619615. Preprint, not peer reviewed.

36. Maria Rigau, David Juan, Alfonso Valencia, and Daniel Rico. Intronic cnvs and gene expression variation in human populations. PLoS Genetics, 15(1):e1007902, 01 2019. doi: 10.1371/journal.pgen.1007902.

37. Sheriar G. Hormuzdi, Risto Penttinen, Rudolf Jaenisch, and Paul Bornstein. A gene-targeting approach identifies a function for the first intron in expression of the α1(i) collagen gene. Molecular and Cellular Biology, 18(6):3368–3375, 06 1998. doi: 10.1128/mcb.18.6.3368.

38. Jin Wang, Xiaoyu Xi, Shifeng Zhao, Xiaolei Wang, Lixia Yao, Jinlin Feng, and Rong Han. Introns in the naa50 gene act as strong enhancers of tissue-specific expression in arabidopsis. Plant Science, 324:111422, 11 2022. doi: 10.1016/j.plantsci.2022.111422.

39. Masahiko Yamaguchi, Yoko Watanabe, Takuji Ohtani, Shin’ichi Takeda, Hiroshi Yamamoto, and So-ichiro Fukada. Calcitonin receptor signaling inhibits muscle stem cells from escaping the quiescent state and the niche. Cell Reports, 13(2):302–314, 10 2015. doi: 10.1016/j.celrep.2015.08.064.

40. Alexander S. Ham, Shuo Lin, Alice Tse, Marco Thürkauf, Timothy J. McGowan, Lena Jörin, Filippo Oliveri, and Markus A. Rüegg. Single-nuclei sequencing of skeletal muscle reveals subsynaptic-specific transcripts involved in neuromuscular junction maintenance. Nature Communications, 16:2220, 03 2025. doi: 10.1038/s41467-025-12017-0.

41. Robert D. Barber, Dan W. Harmer, Robert A. Coleman, and Brian J. Clark. Gapdh as a housekeeping gene: analysis of gapdh mrna expression in a panel of 72 human tissues. Physiological Genomics, 21(3):389–395, 05 2005. doi: 10.1152/physiolgenomics.00025.2005.

42. Jianqiang Bao, Shuiqiao Yuan, Ashley Maestas, Bhupal P. Bhetwal, Andrew Schuster, and Wei Yan. Stk31 is dispensable for embryonic development and spermatogenesis in mice. Molecular Reproduction and Development, 80(10):786, 2013. doi: 10.1002/mrd.22225.

43. V. A. Traag, Ludo Waltman, and Nees Jan van Eck. From louvain to leiden: guaranteeing well-connected communities. Scientific Reports, 9:5233, 03 2019. doi: 10.1038/s41598-019-41695-z.

44. D. Djureinovic, L. Fagerberg, B. Hallström, A. Danielsson, C. Lindskog, M. Uhlén, and F. Pontén. The human testis-specific proteome defined by transcriptomics and antibody-based profiling. Molecular Human Reproduction, 20(6):476–488, 06 2014. doi: 10.1093/molehr/gau018.

45. Uku Raudvere, Liis Kolberg, Ivan Kuzmin, Tambet Arak, Priit Adler, Hedi Peterson, and Jaak Vilo. g:profiler: a web server for functional enrichment analysis and conversions of gene lists (2019 update). Nucleic Acids Research, 47(W1):W191–W198, 07 2019. doi: 10.1093/nar/gkz369.

46. Inna Gitelman. Twist protein in mouse embryogenesis. Developmental Biology, 189(2): 205–214, 09 1997. doi: 10.1006/dbio.1997.8614.

47. Hector L. Franco, José Casasnovas, José R. Rodríguez-Medina, and Carmen L. Cadilla. Redundant or separate entities?—roles of twist1 and twist2 as molecular switches during gene transcription. Nucleic Acids Research, 39(4):1177–1186, 10 2010. doi: 10.1093/nar/gkq890.

48. Jiafa Ren and Steven D. Crowley. Twist1: A double-edged sword in kidney diseases. Kidney Diseases, 6(4):247–257, 01 2020. doi: 10.1159/000505188.

49. Qingfeng Wu, Shiren Sun, Lei Wei, Minna Liu, Hao Liu, Ting Liu, Ying Zhou, Qing Jia, D. Wang, Zhen Yang, Menglu Duan, Xiaoxia Yang, Peisong Gao, and Xiaoxuan Ning. Twist1 regulates macrophage plasticity to promote renal fibrosis through galectin-3. Cellular and Molecular Life Sciences, 79(3):137, 02 2022. doi: 10.1007/s00018-022-04137-0.

50. Lianqin Sun, Lishan Liu, Juanjuan Jiang, Kang Liu, Jingfeng Zhu, Lin Wu, Xiaohan Lu, Zhimin Huang, Yanggang Yuan, Steven D. Crowley, Huijuan Mao, Changying Xing, and Jiafa Ren. Transcription factor twist1 drives fibroblast activation to promote kidney fibrosis via signaling proteins prrx1/tnc. Kidney International, 106(5):840–855, 11 2024. doi: 10.1016/j.kint.2024.07.028.

51. Yan Xu, Yixiang Xu, Lan Liao, Niya Zhou, Sarah M. Theissen, Xin-Hua Liao, Hoang Nguyen, Thomas Ludwig, Li Qin, Jarrod D. Martinez, Jun Jiang, and Jianming Xu. Inducible knock-out of twist1 in young and adult mice prolongs hair growth cycle and has mild effects on general health, supporting twist1 as a preferential cancer target. The American Journal of Pathology, 183(4):1281–1292, 10 2013. doi: 10.1016/j.ajpath.2013.06.021.

52. Jibin Zhang, Jinsoo Ahn, Yeunsu Suh, Seongsoo Hwang, Michael E. Davis, and Kichoon Lee. Identification of CTLA2A, DEFB29, WFDC15B, SERPINA1F and MUP19 as novel tissue-specific secretory factors in mouse. PLOS ONE, 10(5):e0124962, 05 2015. doi: 10.1371/journal.pone.0124962.

53. Xiao Wang, Fanyi Qiu, Junjie Yu, Meiyang Zhou, Anjian Zuo, Xiaojiang Xu, Xiao-Yang Sun, and Zhengpin Wang. Transcriptome profiling of the initial segment and proximal caput of mouse epididymis. Frontiers in Endocrinology, 14:1190890, 05 2023. doi: 10.3389/fendo.2023.1190890.

54. Shuanggang Hu, Guangxin Yao, Xiangqi Li, Zijiang Chen, and Yun Sun. Androgen receptor binding sites identified in mouse testis. Acta Biochimica et Biophysica Sinica, 45(8):795–797, 2013. doi: 10.1093/abbs/gmt076. Advance Access Publication 9 July 2013.

55. Steve Dorus, Elizabeth R. Wasbrough, Jennifer Busby, Elaine C. Wilkin, and Timothy L. Karr. Sperm proteomics reveals intensified selection on mouse sperm membrane and acrosome genes. Molecular Biology and Evolution, 27(6):1235–1246, 06 2010. doi: 10.1093/molbev/msq007.

56. Hui Qian, Xing Deng, Zhao-Wei Huang, Ji Wei, Chen-Hong Ding, Ren-Xin Feng, Xin Zeng, Yue-Xiang Chen, Jin Ding, Lei Qiu, Zhen-Lin Hu, Xin Zhang, Hong-Yang Wang, Jun-Ping Zhang, and Wei-Fen Xie. An hnf1α-regulated feedback circuit modulates hepatic fibrogenesis via the crosstalk between hepatocytes and hepatic stellate cells. Cell Research, 25: 930–945, 07 2015. doi: 10.1038/cr.2015.82.

57. Taro E. Akiyama, Jerrold M. Ward, and Frank J. Gonzalez. Regulation of the liver fatty acid-binding protein gene by hepatocyte nuclear factor 1α (hnf1α): Alterations in fatty acid homeostasis in hnf1α-deficient mice. Journal of Biological Chemistry, 275(35):27117–27122, 09 2000. doi: 10.1016/S0021-9258(19)61487-0.

58. Qi Ni, Kai Ding, Ke-Qi Wang, Jin He, Chuan Yin, Jian Shi, Xin Zhang, Wei-Fen Xie, and Yong-Quan Shi. Deletion of hnf1α in hepatocytes results in fatty liver-related hepatocellular carcinoma in mice. FEBS Letters, 591(11):1481–1491, 05 2017. doi: 10.1002/1873-3468.12689.

59. David Q. Shih, Seamus Screenan, Karla N. Munoz, Lou Philipson, Marco Pontoglio, Moshe Yaniv, Kenneth S. Polonsky, and Markus Stoffel. Loss of hnf-1α function in mice leads to abnormal expression of genes involved in pancreatic islet development and metabolism. Diabetes, 50(11):2472–2480, 11 2001. doi: 10.2337/diabetes.50.11.2472.

60. Marco Pontoglio, Dominique Prié, Claire Cheret, Antonia Doyen, Christine Leroy, Philippe Froguel, Gilberto Velho, Moshe Yaniv, and Gérard Friedlander. Hnf1α controls renal glucose reabsorption in mouse and man. EMBO Reports, 1(4):359–365, 2000. doi: 10.1093/embo-reports/kvd071.

61. Giorgia Benegiamo, Giacomo V. G. von Alvensleben, Sandra Rodríguez-López, Ludger J. E. Goeminne, Alexis M. Bachmann, Jean-David Morel, Ellen Broeckx, Jing Ying Ma, Vinicius Carreira, Sameh A. Youssef, Nabil Azhar, Dermot F. Reilly, Katharine D’Aquino, Shannon Mullican, Maroun Bou-Sleiman, and Johan Auwerx. The genetic background shapes the susceptibility to mitochondrial dysfunction and nash progression. Journal of Experimental Medicine, 220(4):e20221738, 04 2023. doi: 10.1084/jem.20221738.

62. Arthur Liberzon, Chet Birger, Helga Thorvaldsdóttir, Mahmoud Ghandi, Jill P. Mesirov, and Pablo Tamayo. The molecular signatures database hallmark gene set collection. Cell Systems, 1(6):417–425, 12 2015. doi: 10.1016/j.cels.2015.12.004.

63. Gregory R. Keele, Ji-Gang Zhang, John Szpyt, Ron Korstanje, Steven P. Gygi, Gary A. Churchill, and Devin K. Schweppe. Global and tissue-specific aging effects on murine proteomes. Cell Reports, 42(7):112715, 07 2023. doi: 10.1016/j.celrep.2023.112715.

64. Esther M. Speer, Atilade A. Adedeji, Joyce Lin, Alexandra Khorasanchi, Asma Rasheed, Maya Bhat, Kelly Mackenzie, Randolph Hennigar, Kimberly J. Reidy, and Robert P. Woroniecki. Attenuation of acute kidney injury in a murine model of neonatal Escherichia coli sepsis. Frontiers in Cellular and Infection Microbiology, 14:1507914, 2025. doi: 10.3389/fcimb.2024.1507914.

65. Akinobu Ochi, Dong Chen, Wibke Schulte, Lin Leng, Nickolas Moeckel, Marta Piecychna, Luisa Averdunk, Christian Stoppe, Richard Bucala, and Gilbert Moeckel. Mif-2/d-dt enhances proximal tubular cell regeneration through slpi- and atf4-dependent mechanisms. American Journal of Physiology-Renal Physiology, 313(3):F767–F780, 05 2017. doi: 10.1152/ajprenal.00683.2016.

66. Alexander Lex, Nils Gehlenborg, Hendrik Strobelt, Romain Vuillemot, and Hanspeter Pfister. Upset: Visualization of intersecting sets. IEEE Transactions on Visualization and Computer Graphics, 20(12):1983–1992, Dec 2014. ISSN 1941-0506. doi: 10.1109/TVCG.2014.2346248.

67. Sha Li, Chenghai Wang, Xiaxia Zhang, and Wen Su. Cytochrome p450 omega-hydroxylase 4a14 attenuates cholestatic liver fibrosis. Frontiers in Physiology, 12:688259, 05 2021. doi: 10.3389/fphys.2021.688259.

68. Lingyun Xiong, Jing Liu, Seung Yub Han, Junhyong Kim, Adam L. MacLean, and Andrew P. McMahon. Direct androgen receptor control of sexually dimorphic gene expression in the mammalian kidney. Developmental Cell, 58(21):2338–2358.e5, 11 2023. doi: 10.1016/j.devcel.2023.09.019.

69. Baoping Hu, Yuhe Wang, Zhongtao Wang, Xue He, Li Wang, Dongya Yuan, Yongjun He, Tianbo Jin, and Shumei He. Association of slc11a1 polymorphisms with tuberculosis susceptibility in the chinese han population. Frontiers in Genetics, 13:899124, July 2022. doi: 10.3389/fgene.2022.899124.

70. Kazunobu Ouchi, Yoichi Suzuki, Taro Shirakawa, and Fumio Kishi. Polymorphism of slc11a1 (formerly nramp1) gene confers susceptibility to kawasaki disease. Journal of Infectious Diseases, 187(2):326–329, January 2003. doi: 10.1086/345878.

71. Mariam Sophie, Abdul Hameed, Akhtar Muneer, Azam J. Samdani, Saima Saleem, and Abid Azhar. Slc11a1 polymorphisms and host susceptibility to cutaneous leishmaniasis in pakistan. Parasites & Vectors, 10(12):12, January 2017. doi: 10.1186/s13071-016-1954-y.

72. Zahra Kavian, Saman Sargazi, Mahdi Majidpour, Mohammad Sarhadi, Ramin Saravani, Mansour Shahraki, Shekoufeh Mirinejad, Milad Heidari Nia, and Maryam Piri. Association of slc11a1 polymorphisms with anthropometric and biochemical parameters describing type 2 diabetes mellitus. Scientific Reports, 13:6195, April 2023. doi: 10.1038/s41598-023-33469-y.

73. Jenefer M. Blackwell, Tapasree Goswami, Carlton A. W. Evans, Dean Sibthorpe, Natalie Papo, Jacqueline K. White, Susan Searle, E. Nancy Miller, Christopher S. Peacock, Hiba Mohammed, and Muntaser Ibrahim. Slc11a1 (formerly nramp1) and disease resistance. Cellular Microbiology, 3(12):773–784, December 2001. doi: 10.1046/j.1462-5822.2001.00150.x.

74. Jorge Eliécer Mario-Vásquez, Carlos Andrés Naranjo-González, Jehidys Montiel, Lina M. Zuluaga, Ana M. Vásquez, Alberto Tobón-Castaño, Gabriel Bedoya, and Cesar Segura. Association of variants in il1b, tlr9, trem1, il10ra, and cd3g and native american ancestry on malaria susceptibility in colombian populations. Infection, Genetics and Evolution, 87: 104675, January 2021. doi: 10.1016/j.meegid.2020.104675.

75. Dan N. Predescu, Babak Mokhlesi, and Sanda A. Predescu. X-inactive-specific transcript: a long noncoding rna with a complex role in sex differences in human disease. Biology of Sex Differences, 15:101, December 2024. doi: 10.1186/s13293-024-00681-5. Review article.

76. Eli Eisenberg and Erez Y. Levanon. Human housekeeping genes, revisited. Trends in Genetics, 29(10):569–574, 2013. doi: 10.1016/j.tig.2013.05.010.

77. Chintan J. Joshi, Wenfan Ke, Anna Drangowska-Way, Eyleen J. O’Rourke, and Nathan E. Lewis. What are housekeeping genes? PLOS Computational Biology, 18(7):e1010295, July 2022. doi: 10.1371/journal.pcbi.1010295. Version 2, Open Access, Peer-reviewed.

78. Roy D. Dar, Brandon S. Razooky, Abhyudai Singh, and Leor S. Weinberger. Transcriptional burst frequency and burst size are equally modulated across the human genome. Proceedings of the National Academy of Sciences, 109(43):17454–17459, 10 2012. doi: 10.1073/pnas.1213530109.

79. Ido Golding, Johan Paulsson, Scott M. Zawilski, and Edward C. Cox. Real-time kinetics of gene activity in individual bacteria. Cell, 123(6):1025–1036, 12 2005. doi: 10.1016/j.cell.2005.09.031.

80. Martin Rosvall and Carl T. Bergstrom. Mapping change in large networks. PLOS ONE, 5 (1):e8694, January 2010. doi: 10.1371/journal.pone.0008694.

81. Tara Chari and Lior Pachter. The specious art of single-cell genomics. PLOS Computational Biology, 19(8):e1011288, August 2023. doi: 10.1371/journal.pcbi.1011288.

82. Yoav Gilad and Orna Mizrahi-Man. A reanalysis of mouse encode comparative gene expression data. F1000Research, 4:121, 2015. doi: 10.12688/f1000research.6536.1.

83. David Brawand, Magali Soumillon, Anamaria Necsulea, Philippe Julien, Gábor Csárdi, Patrick Harrigan, Manuela Weier, Angélica Liechti, Ayinuer Aximu-Petri, Martin Kircher, Frank W. Albert, Ulrich Zeller, Philipp Khaitovich, Frank Grützner, Sven Bergmann, Rasmus Nielsen, Svante Pääbo, and Henrik Kaessmann. The evolution of gene expression levels in mammalian organs. Nature, 478:343–348, 2011. doi: 10.1038/nature10532.

84. Anamaria Necsulea and Henrik Kaessmann. Evolutionary dynamics of coding and non-coding transcriptomes. Nature Reviews Genetics, 15:734–748, 2014. doi: 10.1038/nrg3802.

85. F. Alexander Wolf, Philipp Angerer, and Fabian J. Theis. Scanpy: large-scale single-cell gene expression data analysis. Genome Biology, 19:15, 2018. doi: 10.1186/s13059-017-1382-0.

86. Tim Stuart, Andrew Butler, Paul Hoffman, Christoph Hafemeister, Efthymia Papalexi, William M. III Mauck, Yuhan Hao, Marlon Stoeckius, Peter Smibert, and Rahul Satija. Comprehensive integration of single-cell data. Cell, 177(7):1888–1902.e21, 2019. doi: 10.1016/j.cell.2019.05.031.

