## Supplementary Figures for "Determining gene specificity from multivariate single-cell RNA sequencing data"

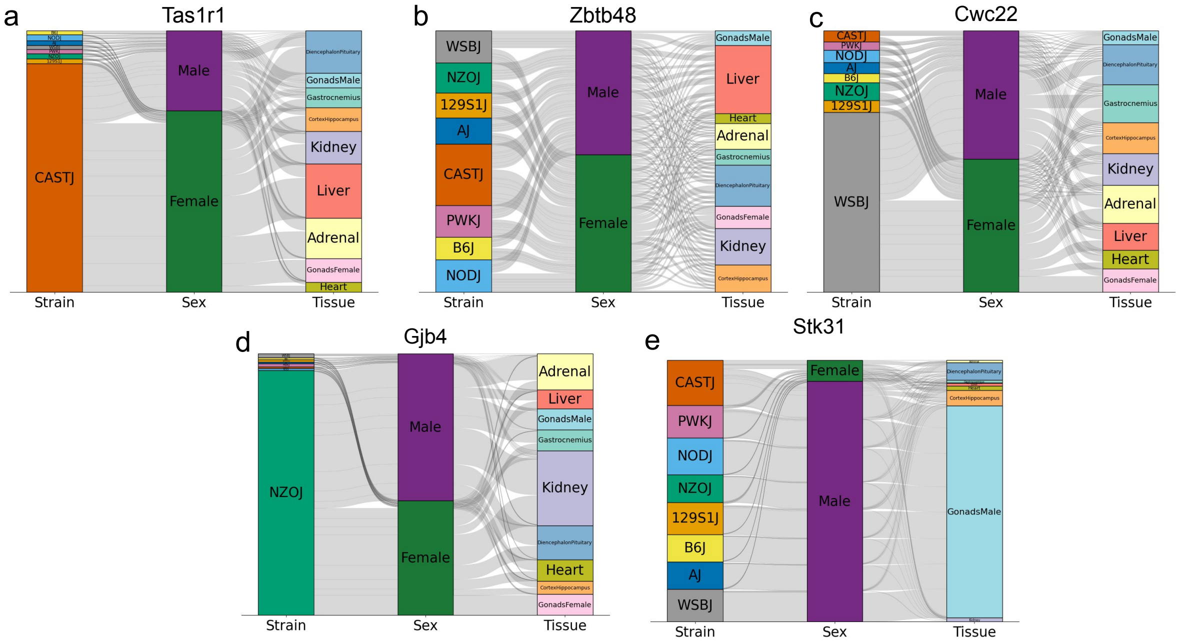


Figure 1: **Alluvial plots showing pseudo-bulked expression across partitions of strain-specific genes a.** *Tas1r1* a CASTJ specific genes. **b.** *Zbtb48*, a gene upstream of *Tas1r1* with slight bias towards CASTJ. **c.** *Cwc22*, a WSBJ specific gene [2]. **d.** *Gjb4*, a NZOJ specific gene [3]. **e.** *Stk31*, a gene that displays strain driven cell type switching in gastrocnemius tissue.

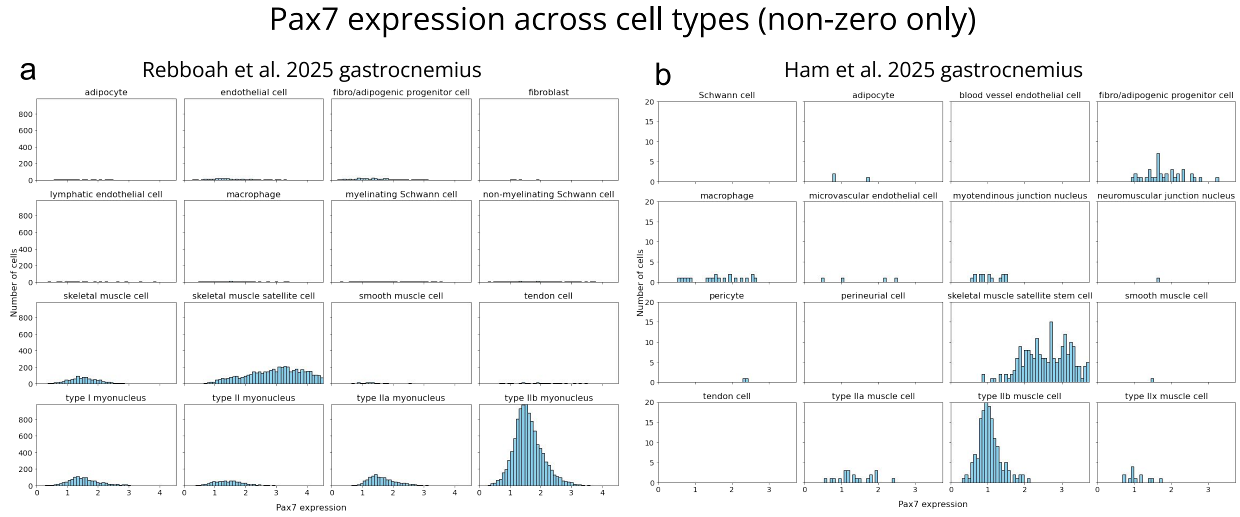


Figure 2: **Pax7 expression across cell types** Histograms of sequence depth and log1p normalized counts across gastrocnemius cells for *Pax7* in **a.** 8cube Rebboah et al. 2026 [5] and **b.** Ham et al. 2025 [4]

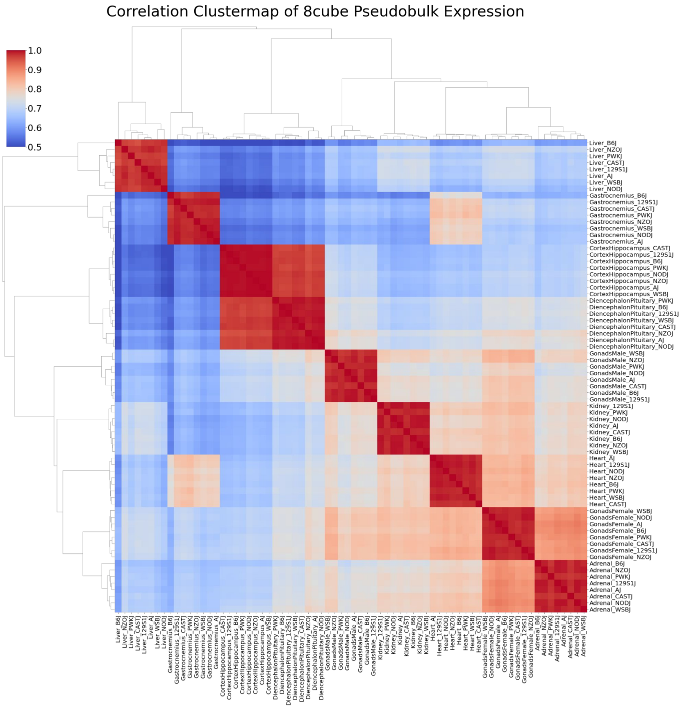


Figure 3: **Correlation cluster map of 8cube data pseudo-bulked by strain and tissue** [5]

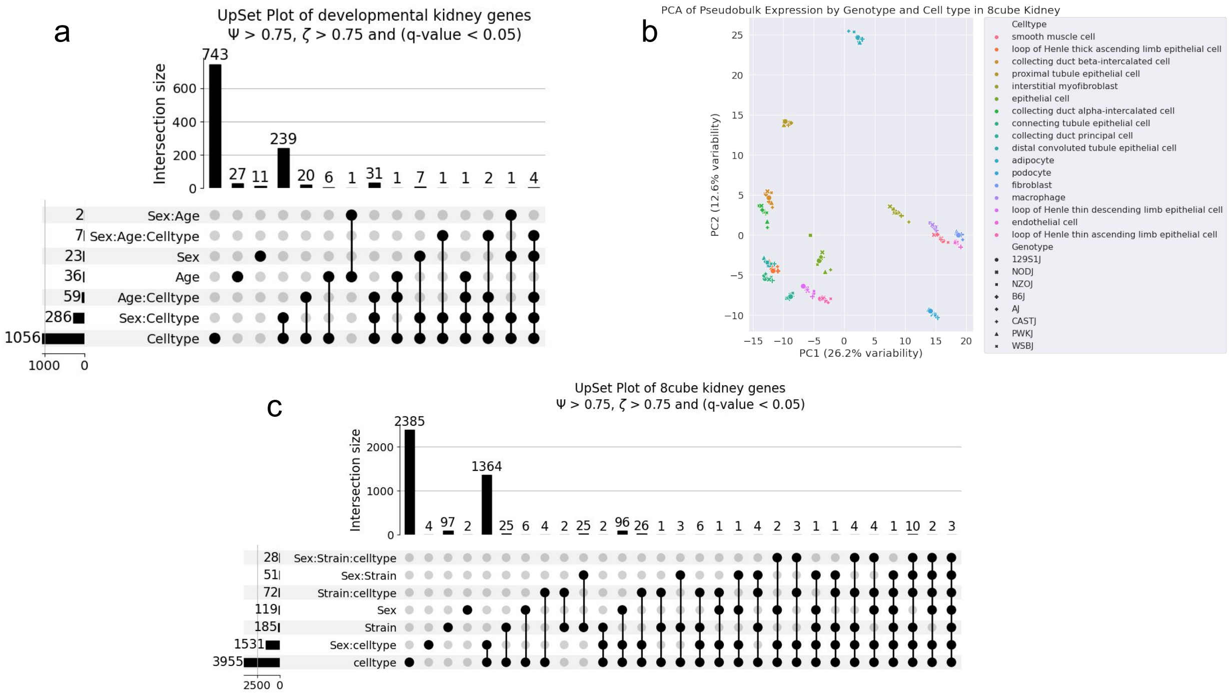


Figure 4: **Specificity trends in the kidney a.** Upset plot of developmental kidney specificity generated from Chen et al. 2025 dataset [1]. We selected highly specific genes partitioned by Sex, Age, Celltype and their 2-way and 3-way interaction terms. Thresholds used for highly specific gene selection are $\Psi > 0.75$ and $\zeta > 0.75$. Global testing correction was performed across all 7 partitioned and genes selected passed a significance threshold of 0.05. **b.** PCA plot of 8cube kidney data pseudo-bulked by cell type (color) and strain(shape) [5]. **c.** Upset plot of 8cube kidney specificity [5]. We selected highly specific genes partitioned by Sex, Strain, Celltype and their 2-way and 3-way interaction terms. Thresholds used for highly specific gene selection are $\Psi > 0.75$ and $\zeta > 0.75$. Global testing correction was performed across all 7 partitioned and genes selected passed a significance threshold of 0.05.

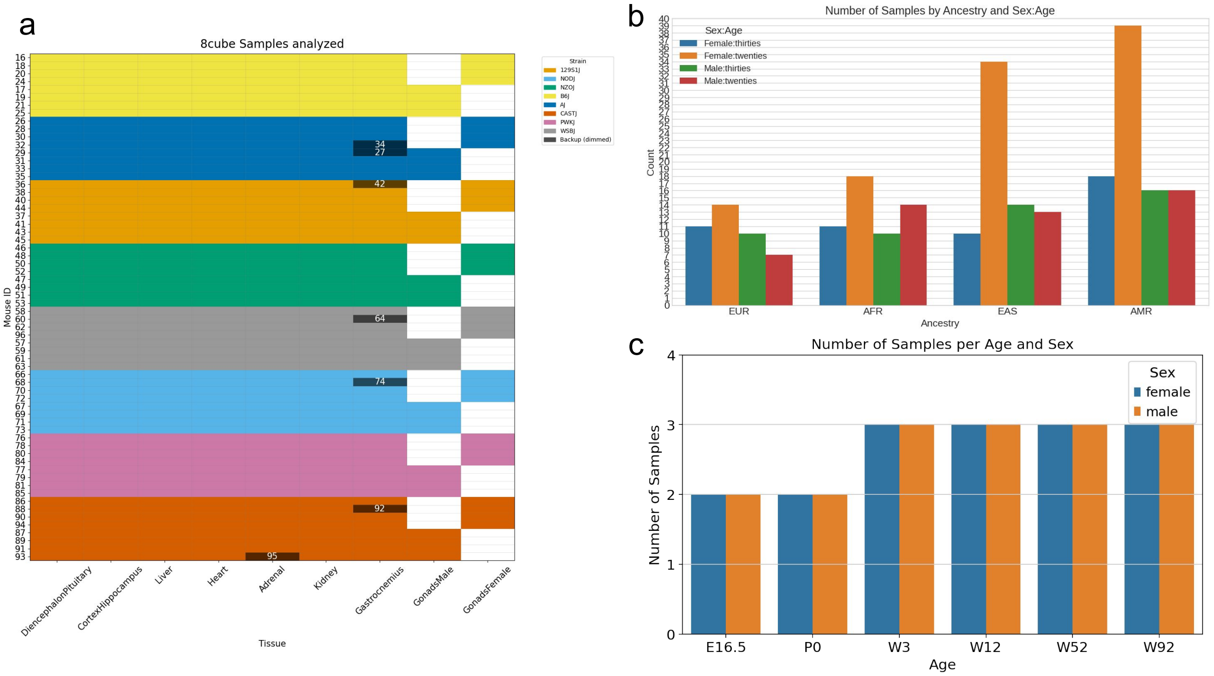


Figure 5: **Biological replicates analyzed from each dataset a.** Samples analyzed from 8cube data, colored by strain. Replacement tissue samples from additional mice shaded in black. **b.** Number of samples analyzed from human PBMCs collected from 255 diverse individuals, grouped by sex, ancestry and age (binned as twenties and thirties). **c.** Number of biological replicates analyzed from developmental kidney data [1], grouped by sex and age.
