## Supplementary Note for "Determining gene specificity from multivariate single-cell RNA sequencing data"

### 1 Entropy basis for the specificity score

Let $A$ be a partition of $n$ cells into groups (e.g. tissues, clusters, or sexes), and let

$$p = (p_1, \dots, p_n) \in \Delta_{n-1}, \qquad \Delta_{n-1} = \{p_i \geq 0 : \sum_i p_i = 1\},$$

be the normalized expression profile of a gene across cells. The distribution $p_A = (p_a)_{a \in A}$ has $p_a = \sum_{i \in a} p_i$. A partition $B$ refines $A$, written $A \prec B$, if every block of $B$ lies within a block of $A$.

We present four axioms for the class of entropy-decomposition scores:

**A1. Regularity and nontriviality.** $I$ is continuous and nonnegative on every finite probability simplex, $I(1) = 0$, and $I(1/2, 1/2) > 0$.

**A2. Symmetry.** $I$ is unchanged by permuting the entries of a probability vector.

**A3. Hierarchical grouping.** For every partition $A$ of the cells, with $p(\cdot \mid a) = (p_i/p_a)_{i \in a}$ for $p_a > 0$,

$$I(p) = I(p_A) + \sum_{a \in A : p_a > 0} p_a I\big(p(\cdot \mid a)\big).$$

**A4. Relative within-group contribution.** For $I(p) > 0$, the score $\Psi(p; A)$ is the fraction of total information in the within-group terms of A3:

$$\Psi(p; A) = \frac{\sum_{a \in A : p_a > 0} p_a I\big(p(\cdot \mid a)\big)}{I(p)}.$$

Faddeev's theorem characterizes $I$ under A1–A3. A4 then fixes $\Psi$ within this class of normalized entropy-decomposition scores. We recall Faddeev's theorem for completeness.

**Theorem 1** (Faddeev, 1956 [1])**.** *Let $I$ be a family of continuous, symmetric, nonnegative functionals on finite probability vectors, with $I(1) = 0$ and $I(1/2, 1/2) > 0$, satisfying the grouping axiom*

$$I(p_{ij}) = I(p_i) + \sum_i p_i \, I(q_{j|i}), \quad \textit{where } p_{ij} = p_i q_{j|i},$$

*and for each $i$ with $p_i > 0$, the conditional distribution $q_{j|i}$ satisfies*

$$q_{j|i} = \frac{p_{ij}}{p_i}, \quad \sum_j q_{j|i} = 1.$$

*Then there exists a constant $\kappa > 0$ such that, for every finite probability vector,*

$$I(p) = \kappa \Big( - \sum_i p_i \log p_i \Big).$$

**Theorem 2** (Characterization of the normalized within-group score)**.** *Under A1–A4, for every profile $p$ with $I(p) > 0$, the score is uniquely determined within this class and equals*

$$\Psi(p; A) = \frac{I(p) - I(p_A)}{I(p)} = 1 - \frac{H(p_A)}{H(p)}, \qquad H(p) = -\sum_i p_i \log p_i.$$

*The value is independent of the multiplicative normalization of $I$. The theorem makes no uniqueness claim among specificity measures outside A1–A4.*

*Proof.* By A1–A3 and Theorem 1, $I = \kappa H$ for some $\kappa > 0$. A3 and A4 give

$$\Psi(p; A) = \frac{I(p) - I(p_A)}{I(p)}.$$

Substituting $I = \kappa H$ cancels $\kappa$. The Shannon chain rule gives

$$H(p) - H(p_A) = \sum_{a \in A} p_a H\big(p(\cdot \mid a)\big),$$

so the numerator is the weighted within-group entropy. □

This score lies in $[0, 1]$ whenever $H(p) > 0$. It equals 1 if all expression is found in a single block and equals 0 if each occupied block contains at most one cell with positive expression. Uniform expression across all cells generally does not give 0: for $r$ equal blocks of $m$ cells, it gives $1 - \log r / \log(rm)$. The score is invariant under relabeling cells within blocks or relabeling blocks and is continuous wherever $H(p) > 0$. In general, it has no continuous extension to point-mass profiles, where its denominator vanishes.

To simplify the discussion in what follows, for a distribution $p = (p_1, \ldots, p_n)$ and a partition $A$ of $[n]$ into groups with coarse law $p_A$, we use $E_T(p)$ to denote the total Shannon entropy $H(p)$, $E_B(p; A)$ to denote the between-group entropy $H(p_A)$ and $E_W(p; A)$ to denote the within-group entropy

$$E_W(p; A) := E_T(p) - E_B(p; A).$$

The normalized within-group score at level $A$ is then given by

$$\Psi(p; A) := \frac{E_W(p; A)}{E_T(p)} = 1 - \frac{E_B(p; A)}{E_T(p)}.$$

### 2 Additivity

The additive property belongs to the underlying entropy. For partitions $A \prec B$, write $H(B \mid A) = \sum_{a \in A : p_a > 0} p_a H((p_b/p_a)_{b \in B,\, b \subseteq a})$ for the entropy of the subblocks of $B$ conditional on their blocks in $A$. The Shannon chain rule gives $H(p_B) = H(p_A) + H(B \mid A)$, and thus

$$E_W(p; A) = E_W(p; B) + H(B \mid A), \qquad \Psi(p; A) = \Psi(p; B) + \frac{H(B \mid A)}{H(p)}. \tag{1}$$

The latter identity holds when $H(p) > 0$; the normalized scores themselves do not obey the unweighted Shannon chain rule. Suppose that we have two tissues (A and B), each containing two cell types. Let the joint probability vector for a gene's expression be

$$p = (p_{\mathrm{A},1},\, p_{\mathrm{A},2},\, p_{\mathrm{B},1},\, p_{\mathrm{B},2}).$$

Summing within tissues gives $p_{\text{tissue}} = (p_{\text{A}},\, p_{\text{B}})$, where $p_{\text{A}} = p_{\text{A},1} + p_{\text{A},2}$ and $p_{\text{B}} = p_{\text{B},1} + p_{\text{B},2}$. Conditional distributions within tissues define $q_{\text{cell}|\text{A}} = \big(p_{\text{A},1}/p_{\text{A}},\, p_{\text{A},2}/p_{\text{A}}\big)$ and $q_{\text{cell}|\text{B}} = \big(p_{\text{B},1}/p_{\text{B}},\, p_{\text{B},2}/p_{\text{B}}\big)$.

Here $H(B \mid A)$ is the weighted average of the conditional entropies in the two tissues, with weights $p_{\text{A}}$ and $p_{\text{B}}$. Equation (1) relates the tissue-level score to the score after refinement by cell type.

**Example.** Consider the gene count table:

| | cell 1 | cell 2 | total (tissue) |
|---|---|---|---|
| A | 40 | 10 | 50 |
| B | 25 | 15 | 40 |
| Total | | | 90 |

so that

$$p = \left(\tfrac{40}{90},\, \tfrac{10}{90},\, \tfrac{25}{90},\, \tfrac{15}{90}\right), \qquad p_{\text{tissue}} = \left(\tfrac{50}{90},\, \tfrac{40}{90}\right).$$

The conditional distributions within tissues are

$$q_{\text{cell}|\text{A}} = (0.8,\, 0.2), \qquad q_{\text{cell}|\text{B}} = (0.625,\, 0.375).$$

Using base-2 Shannon entropy,

$$H(p) = -\sum_i p_i \log_2 p_i,$$

we obtain

$$H(p) = 1.8163, \qquad H(p_{\text{tissue}}) = 0.9911.$$

The within-tissue entropy is

$$H_{\text{within}} := \tfrac{50}{90}\, H(q_{\text{cell}|\text{A}}) + \tfrac{40}{90}\, H(q_{\text{cell}|\text{B}}),$$

where

$$H(q_{\text{cell}|\text{A}}) = 0.7219, \qquad H(q_{\text{cell}|\text{B}}) = 0.9544,$$

so that

$$H_{\text{within}} = 0.8253.$$

These values satisfy the Shannon chain rule

$$H(p) = H(p_{\text{tissue}}) + H_{\text{within}} = 0.9911 + 0.8253 = 1.8163.$$

Moreover,

$$E_T(p) = H(p) = 1.8163,$$
$$E_B(p;\text{tissue}) = H(p_{\text{tissue}}) = 0.9911,$$
$$E_W(p;\text{tissue}) = H_{\text{within}} = 0.8253.$$

The specificity at the tissue level is

$$\Psi_{\text{tissue}} = \Psi(p;\text{tissue}) = \frac{E_W(p;\text{tissue})}{E_T(p)} = 1 - \frac{E_B(p;\text{tissue})}{E_T(p)} = 1 - \frac{0.9911}{1.8163} = 0.4544.$$

Equivalently, the between-tissue contribution is

$$\frac{E_B(p; \text{tissue})}{E_T(p)} = \frac{0.9911}{1.8163} = 0.5456,$$

and the two normalized components sum to one:

$$0.4544 + 0.5456 = 1.$$

Thus, the normalized within-tissue and between-tissue entropy contributions are complementary, with no dependence on the logarithm base. If the four cells are the blocks of the fine partition $B$, then $\Psi(p; B) = 0$ and $H(B \mid \text{tissue})/H(p) = 0.4544$, in agreement with Equation (1).

### 3 Order invariance and hierarchical decomposition

The score for a fixed final partition is independent of the order used to reach that partition. To illustrate, let $X$, $Y$, and $Z$ denote strain, tissue, and sex, and let $J$ be their joint partition of the same cells. The Shannon chain rule gives

$$\begin{aligned} H(p_J) &= H(X) + H(Y \mid X) + H(Z \mid X, Y) \\ &= H(Y) + H(Z \mid Y) + H(X \mid Y, Z), \end{aligned} \tag{2}$$

and therefore, when $H(p) > 0$,

$$\Psi(p; J) = 1 - \frac{H(X) + H(Y \mid X) + H(Z \mid X, Y)}{H(p)} = 1 - \frac{H(Y) + H(Z \mid Y) + H(X \mid Y, Z)}{H(p)}.$$

The individual conditional contributions can change with the order; their sum, and hence the score for the same joint partition, cannot. Equation (1) gives the corresponding within-entropy relationship at each refinement step.

### 4 Five-pattern simulation benchmark

#### 4.1 Benchmark design

We next evaluated whether gene-ranking methods recover five known single-cell specificity patterns from a simulated 3,000-gene dataset. The simulation contained two strains, StrainA and StrainB, and four cell types, CT1–CT4. By default, each strain–cell-type block contained 500 cells, split evenly across four mice per strain. Counts were sampled from a gamma–Poisson model with cell-depth and mouse-level nuisance effects.

Genes 0–999 contained the planted signal. Each planted category contained 200 genes, and genes 1000–2999 were null/background genes. For the run summarized here, the target fold-change was 8×, the dispersion was 8.0, and the random seed was 123. Raw counts were saved in the `raw_counts` layer; the primary AnnData matrix was depth-normalized to a target sum of 10,000 counts per cell and `log1p`-transformed for ember.

Table 1: Planted truth categories in the simulation.

| Category | Gene range | Target blocks | Biological pattern |
|---|---|---|---|
| 1 CT | gene_0–gene_199 | CT1 | CT1-specific in both strains |
| 2 CTs | gene_200–gene_399 | CT1; CT2 | CT1/CT2-specific in both strains |
| Housekeeping | gene_400–gene_599 | all | High expression in all cells |
| 1 CT, 1 strain | gene_600–gene_799 | StrainA:CT1 | CT1-specific only in StrainA |
| Strain switch | gene_800–gene_999 | StrainA:CT1; StrainB:CT2 | Different target cell types by strain |

Table 2: Partition, ranking statistic, and cutoffs used for each benchmark method and category.

| Method | Categories | Partition | Ranking and cutoffs |
|---|---|---|---|
| ember | 1 CT, 2 CTs, Housekeeping | cell_type | Eligible if $\Psi \geq 0.5$. For 1 CT and 2 CTs, also require a $\Psi$ p-value $< 0.05$ and q-value $< 0.05$; rank by CT1 $\Psi$-block for 1 CT and by $\min(\mathrm{CT1}, \mathrm{CT2})$ $\Psi$-block for 2 CTs. Housekeeping genes are ranked by lowest $\zeta$ among genes with mean raw count $\geq 1.0$; p/q significance is not required for housekeeping. |
| ember | 1 CT, 1 strain; Strain switch | strain_celltype | Eligible if $\Psi \geq 0.5$, the $\Psi$ p-value $< 0.05$, and q-value $< 0.05$. Rank by StrainA:CT1 $\Psi$-block for 1 CT, 1 strain and by the smaller of the StrainA:CT1 and StrainB:CT2 $\Psi$-block values for the strain-switch pattern. |
| DESeq2 [5] | 1 CT, 2 CTs, 1 CT, 1 strain, Strain switch | pseudobulk mouse $\times$ cell_type | Pseudobulk raw counts are modeled with design $\sim$ strain $*$ cell_type. Genes are eligible at adjusted p-value $< 0.05$ and ranked by descending test statistic for the contrast corresponding to the planted pattern. |

| Method | Categories | Partition | Ranking and cutoffs |
| --- | --- | --- | --- |
| DESeq2 [5] | Housekeeping | pseudobulk mouse × cell_type | Genes are treated as housekeeping candidates if the DESeq2 test is not significant (adjusted p-value $\geq 0.05$), then ranked by descending baseMean. |
| variance Partition [3] | 1 CT, 2 CTs | pseudobulk mouse × cell_type | Rank genes by descending fraction of expression variance attributed to `cell_type`. |
| variance Partition [3] | 1 CT, 1 strain; Strain switch | pseudobulk mouse × cell_type | Rank genes by descending fraction of expression variance attributed to `strain_celltype`. |
| variance Partition [3] | Housekeeping | pseudobulk mouse × cell_type | Eligible if summed biological variance from `strain`, `cell_type`, and `strain_celltype` is $\leq 0.05$; rank by descending mean logCPM. |
| Tau [6] | 1 CT, 2 CTs, Housekeeping | cell_type | Compute Tau on mean expression by cell type. Non-housekeeping categories are ranked by descending Tau. Housekeeping is ranked by ascending Tau after requiring mean raw count $\geq 1.0$. |
| Tau [6] | 1 CT, 1 strain; Strain switch | strain_celltype | Compute Tau on mean expression by strain–cell-type block; rank by descending Tau. |

| Method | Categories | Partition | Ranking and cutoffs |
|---|---|---|---|
| PEM [4] | 1 CT, 2 CTs, Housekeeping | cell_type | Compute PEM from summed expression by cell type. Rank by CT1 PEM for 1 CT and by $\min(\text{CT1}, \text{CT2})$ PEM for 2 CTs. Housekeeping is ranked by ascending maximum absolute PEM after requiring mean raw count $\geq 1.0$. |
| PEM [4] | 1 CT, 1 strain; Strain switch | strain_celltype | Compute PEM from summed expression by strain–cell-type block. Rank by StrainA:CT1 PEM for 1 CT, 1 strain and by the smaller of the StrainA:CT1 and StrainB:CT2 PEM values for the strain-switch pattern. |
| Gini [2] | 1 CT, 2 CTs, Housekeeping | cell_type | Compute Gini on mean expression by cell type. Non-housekeeping categories are ranked by descending Gini. Housekeeping is ranked by ascending Gini after requiring mean raw count $\geq 1.0$. |
| Gini [2] | 1 CT, 1 strain; Strain switch | strain_celltype | Compute Gini on mean expression by strain–cell-type block; rank by descending Gini. |

#### 4.2 Recovery results

Table 3: Top-200 recovery for run `simulation_20260720_175005`. Entries are recovered planted genes out of 200.

| Method | 1 CT | 2 CTs | Housekeeping | 1 CT, 1 strain | Strain switch |
|---|---|---|---|---|---|
| ember | 200 | 199 | 200 | 200 | 200 |
| DESeq2 | 183 | 200 | 6 | 83 | 183 |
| variancePartition | 40 | 160 | 7 | 10 | 147 |
| Tau | 200 | 0 | 200 | 199 | 1 |
| PEM | 200 | 200 | 200 | 200 | 200 |
| Gini | 159 | 41 | 200 | 0 | 100 |

Ember and PEM successfully recovered all five planted specificity patterns and distinguished among patterns that shared expression in CT1 but differed in their broader multivariate structure. In contrast, DESeq2 and variancePartition could not reliably distinguish among the four specificity patterns involving CT1, because differential or variable expression is not necessarily specific expression. Both methods also struggled to identify housekeeping genes, for which high, uniform expression—not differential or variable expression—is the defining property. Tau performed well for genes concentrated in a single block but could not recover patterns specific to multiple blocks within the same partition. Gini effectively identified housekeeping genes by measuring expression inequality, but it could not distinguish among more complex multivariate specificity patterns.

### Single-cell versus pseudobulk resolution benchmark

Although ember and PEM recovered all five specificity patterns in the initial benchmark, PEM and the other comparison methods reduce single-cell measurements to aggregated expression summaries. DESeq2 and variancePartition explicitly operate on mouse-level pseudobulks, PEM uses summed expression within each block, and Tau and Gini are calculated from block-level mean expression. These approaches therefore discard information about how expression is distributed among individual cells within a block. In contrast, ember is designed for single-cell data and calculates specificity from the complete cell-level expression distribution.

We therefore designed a resolution challenge to test whether each method could distinguish coherent single-cell specificity from a pseudobulk-identical decoy. Starting from the same simulated dataset, genes gene_1000–gene_1199 were overwritten as paired decoys for the strain-switch genes gene_800–gene_999. For each pair, the decoy gene had exactly the same total raw count as its corresponding true gene in every mouse × strain–cell-type pseudobulk. Within the target blocks StrainA:CT1 and StrainB:CT2, however, the decoy expression was concentrated in only 10% of cells. The coherent and decoy genes were therefore identical after pseudobulking but differed substantially in their cell-level expression distributions.

Each method was evaluated using its top-200 list for the strain-switch pattern. The selected genes were classified as coherent strain-switch genes, rare-cell decoys, or other incorrect genes. The wrong-gene proportion was defined as (decoys + other)/200.

Table 4: Single-cell versus pseudobulk resolution benchmark. Counts describe the composition of each method's top-200 list, and the coherent proportion is the fraction of selected genes exhibiting coherent single-cell expression.

| Method | Coherent | Decoy | Other | Coherent proportion |
|---|---|---|---|---|
| ember | 200 | 0 | 0 | 1.00 |
| DESeq2 | 100 | 100 | 0 | 0.50 |
| variancePartition | 86 | 87 | 27 | 0.43 |
| Tau | 0 | 0 | 200 | 0.00 |
| PEM | 100 | 100 | 0 | 0.50 |
| Gini | 64 | 65 | 71 | 0.32 |

Because the coherent genes and paired decoys had identical pseudobulk expression, PEM and DESeq2 selected them in equal proportions. variancePartition and Gini similarly failed to preferentially recover the coherent genes, while Tau did not recover the multiblock strain-switch pattern. Ember recovered all 200 coherent genes and excluded every rare-cell decoy. Thus, retaining single-cell resolution allowed ember to distinguish genuinely coherent specificity from expression patterns that appeared equally specific only after aggregation.

### 5 Input sensitivity of ember scores

The benchmarks above hold the input matrix, the cell-type labels and the block sizes fixed. We next varied each of these three inputs in turn, using the same simulated dataset, to characterize how much of the recovery reported above depends on choices made before ember is run. All three conditions were scored using the same selection criteria and ranking statistics already described for ember, and recovery is reported against the same five planted truth categories. Recovery is the number of planted genes appearing in ember's top-200 ranked list for that category, as in the benchmarks above, so a gene can be missed either because it failed the selection criteria or because it was outranked.

#### 5.1 Annotation resolution

This benchmark was designed to display how successful detection of specificity changes when partitions are refined. A core property of ember due to its dependence on shannon entropy is it's ability to preserve hierarchical disposability. However, this is limited by the reliability of annotations of a given partition. The more reliable the annotation, the greater the recovered signal. While ember is robust to some amounts of noise, if the annotations are misaligned or not fine enough to capture true biological signal, Psi and Zeta estimates paired with p-values from permutation test will return insignificant specificity results. The below benchmark showcases this limitation of the ember framework. Unknown partition annotations cannot be recovered post-hoc.

The same 4,000 cells and the same expression matrix were scored twice: once with the four coarse `cell_type` labels and once with the eight fine `strain_celltype` labels. The fine labels subdivide each coarse label by strain, so the two partitions are nested and no cells were reclustered or relabelled. For the coarse partition, strain-specific target blocks were mapped to their parent cell types; for the fine partition, coarse targets were scored as the sum of both strain-child block shares.

Table 5: Recovery under coarse and fine partitions of the same cells. Categories are named by their target blocks, as in Table 1. Entries are recovered genes from 200 planted genes.

| Category | Coarse: 4 cell types | Fine: 8 strain × cell types |
|---|---|---|
| CT1 | 195 | 189 |
| CT1 + CT2 | 188 | 180 |
| Housekeeping | 200 | 200 |
| StrainA:CT1 | 5 | 195 |
| StrainA:CT1 + StrainB:CT2 | 11 | 200 |

As expected when a coarse partition is used to recover specificity to finer unknown partitions, ember struggles to recover the meaningful genes. It should be noted that this is an example to demonstrate the effect of refinement of nested partitions, the fine labels used here are simulated strain subdivisions rather than annotated subtypes. Counts describe the effect of nested group definitions and not of clustering resolution.

#### 5.2 Unequal cell numbers

A common concern with single cell data is how differences in sequencing depth of certain cell type populations effects downstream biological analyses. To demonstrate how the ember framework handles such differences we constructed the following experiment: Cell numbers per strain and cell type were varied while the total was held fixed at 4,000 cells. Each base expression profile was repeated by an integer factor within its cell type, so that the empirical expression distribution within every mouse and block is exactly preserved and only abundance changes. Per-strain counts for CT1–CT4 were 500/500/500/500 (balanced), 300/400/600/700 (mild) and 100/200/500/1200 (strong). Categories 1–3 were scored on `cell_type` and categories 4–5 on `strain_celltype`, as in the benchmark above.

Table 6: Recovery under unequal cell numbers with the full selection criteria applied. Categories are named by their target blocks, as in Table 1. Entries are recovered planted genes out of 200. Under strong imbalance no gene in the four block-specific categories passed the significance requirement, so none could be recovered.

| Category | Balanced | Mild imbalance | Strong imbalance |
|---|---|---|---|
| CT1 | 195 | 105 | 0 |
| CT1 + CT2 | 188 | 135 | 0 |
| Housekeeping | 200 | 186 | 200 |
| StrainA:CT1 | 195 | 155 | 0 |
| StrainA:CT1 + StrainB:CT2 | 200 | 198 | 0 |

With the full criteria applied, recovery fell as imbalance increased for the four block-specific categories, and the loss lay entirely in the significance requirement. All 200 planted genes in every category satisfied $\Psi \geq 0.5$ in all three conditions, while the number also passing $p < 0.05$ and $q < 0.05$ fell from 200 in the balanced condition to 105 (CT1), 135 (CT1 + CT2), 163 (StrainA:CT1) and 199 (StrainA:CT1 + StrainB:CT2) under mild imbalance, and to zero in all four under strong imbalance. Housekeeping genes, which carry no significance requirement, were recovered almost completely throughout, at 200, 186 and 200 of 200.

Table 7: Recovery with the significance requirement removed. The ranking statistic, the $\Psi$ cutoff and the housekeeping expression cutoff are unchanged; only the $p$- and $q$-value requirement is dropped. Entries are recovered planted genes out of 200. These counts are descriptive ranking recovery and are not significance-controlled.

| Category | Balanced | Mild imbalance | Strong imbalance |
|---|---|---|---|
| CT1 | 195 | 196 | 177 |
| CT1 + CT2 | 188 | 191 | 192 |
| Housekeeping | 200 | 186 | 200 |
| StrainA:CT1 | 195 | 172 | 104 |
| StrainA:CT1 + StrainB:CT2 | 200 | 198 | 153 |

Removing only the significance requirement leaves recovery essentially unchanged for the two cell-type categories, at 177 and 192 of 200 under strong imbalance against 195 and 188 when balanced, and reduces it for the two strain-resolved categories, to 104 and 153 of 200. The corresponding target blocks differ in size by a factor of five across the conditions: the CT1 block contains 1,000, 600 and 200 cells, and the StrainA:CT1 block 500, 300 and 100. Because $\Psi$ and $\psi_{block}$ are Shannon entropies of a gene's expression shares across cells, they describe the shape of that gene's distribution over blocks rather than the number of cells the blocks contain, and provided a block holds enough cells for its within-block entropy to be estimated stably the ordering of genes by specificity is largely preserved when block sizes change. The score values themselves are not invariant to block size, as in the preprocessing condition below, but the ranking they induce is reasonably stable. The limitation is therefore in the permutation test rather than in the statistic: as a target block shrinks, the observed value is harder to separate from its label-permuted null, and a gene that is still ranked correctly can fail to reach significance. Non-significance under these conditions reflects that loss of power and is not evidence that a gene lacks specificity.

##### 5.3 Expression preprocessing

Single-cell RNA-seq counts are normalized before analysis to reduce technical variation between cells and to make them directly comparable, and the choice of normalization is left to the analyst. We therefore scored the same simulation under three of the choices in common use. Starting from the `raw_counts` layer of the same balanced simulation, the identical cells, genes and labels were scored three times: on raw counts, after depth normalization to a target sum of 10,000 counts per cell, and after a further `log1p` transform. ember performs neither preprocessing step internally, and its probabilities are formed per gene across cells, so a per-cell scaling factor does not cancel and the score values themselves are not invariant to these transformations.

Table 8: Recovery under three expression preprocessing choices. Categories are named by their target blocks, as in Table 1. Entries are recovered planted genes out of 200.

| Category | Raw counts | Library normalized | `log1p` normalized |
|---|---|---|---|
| CT1 | 199 | 200 | 195 |
| CT1 + CT2 | 176 | 180 | 188 |
| Housekeeping | 200 | 200 | 200 |
| StrainA:CT1 | 200 | 199 | 195 |
| StrainA:CT1 + StrainB:CT2 | 200 | 200 | 200 |

Recovery was stable across all three inputs,varying by at most 12 genes out of 200 in any category, and every planted gene passed the selection criteria in all three conditions. The ranked gene sets returned by ember are therefore reasonably stable across the preprocessing choices in common use, even though the underlying score values are not. Raw and depth-normalized values are mathematically admissible inputs to the entropy calculation, but the ember interface recommends depth-normalized, `log1p`-transformed input, and this condition measures input sensitivity rather than endorsing raw counts as an input.

### References

[1] Dmitrii Konstantinovich Faddeev. On the concept of entropy of a finite probabilistic scheme. *Uspekhi Matematicheskikh Nauk*, 11(1):227–231, 1956.

[2] Corrado Gini. *Variabilità e mutabilità: Contributo allo studio delle distribuzioni e delle relazioni statistiche*. Tipografia di Paolo Cuppini, Bologna, Italy, 1912.

[3] Gabriel E. Hoffman and Eric E. Schadt. variancepartition: interpreting drivers of variation in complex gene expression studies. *BMC Bioinformatics*, 17(1):483, 2016.

[4] Lukasz Huminiecki, Andrew T. Lloyd, and Kenneth H. Wolfe. Congruence of tissue expression profiles from gene expression atlas, sagemap and tissueinfo databases. *BMC Genomics*, 4:31, 2003.

[5] Michael I. Love, Wolfgang Huber, and Simon Anders. Moderated estimation of fold change and dispersion for rna-seq data with deseq2. *Genome Biology*, 15:550, 2014.

[6] Itai Yanai, Hila Benjamin, Michael Shmoish, Vered Chalifa-Caspi, Maxim Shklar, Ron Ophir, Arren Bar-Even, Shirley Horn-Saban, Marilyn Safran, Eytan Domany, Doron Lancet, and Orit Shmueli. Genome-wide midrange transcription profiles reveal expression level relationships in human tissue specification. *Bioinformatics*, 21(5):650–659, 2005. All data available at http://genecards.weizmann.ac.il/genenote/; GEO accession GSE803.
